# *Drosophila melanogaster Nepl15* gene deletion promotes anti-aging and anti-obesity phenotypes in a sex-dependent manner

**DOI:** 10.64898/2026.02.20.707128

**Authors:** Shahira Helal Arzoo, Chase Joshua Drucker, Nicolas Jones, Elena Gracheva, Abby Matt, Raymond Hsin, Fei Wang, Chao Zhou, Surya Jyoti Banerjee

## Abstract

Aging and obesity are characterized by comorbidities like declines in fertility, lifespan, gut barrier integrity, cardiac function, and motor activity, and an increase in oxidative stress due to altered nutrient and energy homeostasis. A study on *Drosophila* Neprilysin-like 15 (*Nepl15*) demonstrated that loss of *Nepl15* gene significantly reduced glycogen and glycerolipid reserves in adult males and increased glycogen storage in adult females despite similar food consumption as controls. Therefore, we investigated the sex- and age-specific consequences of *Nepl15* loss on cellular and physiological parameters associated with aging and obesity. We observed that egg production, rate of pupariation, and rate of adult fly eclosion were slightly better in the mutant flies. Interestingly, mutant females, but not males, exhibited significantly extended lifespan. Both sexes demonstrated improved locomotor performance, exercise endurance, gut barrier integrity, and preserved heart rate during progressive aging. At the cellular level, female mutants displayed reduced oxidative stress, elevated *Sod2* expression, and increased ATP levels, all indicative of enhanced cellular health. Mutant males exhibited an increased mitochondrial membrane potential, indicating an enhanced capacity for rapid ATP production in response to enforced activity. Consistently, the energy-sensing kinase *AMPK* expression was reduced in the mutants. The lifespan extension of mutant females was supported by downregulation of *mTOR* and upregulation of *Sirt6* expression. However, in mutant males, both *mTOR* and *Sirt6* were downregulated, potentially contributing to improved physiological health without changing their lifespan. Collectively, our findings establish *Nepl15* knockout mutation promotes anti-aging and anti-obesity health benefits, with stronger effects in female flies.

## Introduction

Obesity is classically attributed to excessive storage of lipids and carbohydrates due to an imbalance between energy intake and expenditure (Dhurandhar, Petersen, & Webster, 2021; Pang, Xie, Chen, & Hu, 2014). Dysregulation of nutrient and energy homeostasis has systemic consequences, including reduced reproductive fitness (Della Torre, Benedusi, Fontana, & Maggi, 2014; Leisegang, Henkel, & Agarwal, 2019; Pereira, Crisóstomo, Sousa, Oliveira, & Alves, 2020), impaired locomotor performance (Collins et al., 2018), compromised cardiac function (Csige et al., 2018), loss of gut barrier integrity (Forlano et al., 2024), and decreased lifespan (Fontaine, Redden, Wang, Westfall, & Allison, 2003; Tam, Morais, & Santosa, 2020); phenotypes that are also widely recognized as hallmarks of aging (Ghosh, Sinha, & Raghunath, 2019; Jura & Kozak, 2016). At the molecular level, these processes are primarily regulated by conserved nutrient-sensing pathways, notably the insulin/ mechanistic Target of Rapamycin (ISS/ *mTOR*) signaling and Sirtuin6 (*Sirt6*)-mediated pathways (González & Hall, 2017; Oldham, 2011; Taylor et al., 2022). mTOR signaling promotes anabolic metabolism, including lipid and glycogen synthesis (Laplante & Sabatini, 2009; Saxton & Sabatini, 2017), and its downregulation has been consistently associated with lifespan extension across species (Bonawitz, Chatenay-Lapointe, Pan, & Shadel, 2007; Evans, Kapahi, Hsueh, & Kockel, 2011). In parallel, *Sirt6* regulates metabolic homeostasis, genomic stability, and oxidative stress responses, thereby influencing stress resistance and longevity (Korotkov, Seluanov, & Gorbunova, 2021; Li et al., 2021; Taylor et al., 2022; Zhou, Tang, & Chen, 2018). Perturbations in these pathways are strongly linked to obesity, metabolic syndrome, and age-associated decline, establishing them as central integrators of nutrient status and physiological output (Zhou et al., 2018).

Mammalian Neprilysin (*Nep*) metalloectopeptidase has been implicated in metabolic and cardiovascular regulation (AlAnazi et al., 2023; Bayes-Genis, Barallat, & Richards, 2016; Buhr, Schiemann, & Meyer, 2023; Nalivaeva, Zhuravin, & Turner, 2020; Schiemann et al., 2022; Standeven et al., 2011). In mammals, neprilysin cleaves substrates such as natriuretic peptides, angiotensin, and insulin and glucagon hormones, thereby modulating pathways involved in metabolism, inflammation, and vascular function (Nalivaeva et al., 2020; Standeven et al., 2011). In obesity, adipocytes become a significant source of Nep; elevated plasma Nep correlates with insulin resistance and metabolic syndrome severity, suggesting Nep as a biomarker and mediator of adipose-driven metabolic dysfunction (Standeven et al., 2011). Yet the broader connection between neprilysin-controlled nutrient homeostasis and metabolic outcomes is incompletely defined.

The *Drosophila* (fruit fly) model has been used to understand the mechanism of nutrient metabolism and energy homeostasis, and related disorders like obesity, diabetes, aging, and lifespan in humans, as both species contain highly conserved cellular pathways, including the ISS/ mTOR and Sirt6, and analogous organs (Chatterjee & Perrimon, 2021; Gáliková & Klepsatel, 2018; Musselman & Kühnlein, 2018; Padmanabha & Baker, 2014; Rajan & Perrimon, 2013). Within the *Drosophila* neprilysin family, Neprilysin-like 15 (*Nepl15*) (Bland, Pinney, Thomas, Turner, & Isaac, 2008; Sitnik et al., 2014) has emerged as a candidate regulator of carbohydrate and lipid storage (S. Banerjee et al., 2021). Unlike typical Neprilysin, *Nepl15* is predicted to be catalytically inactive, suggesting a non-enzymatic regulatory role (S. Banerjee et al., 2021). Expression profiling indicates that *Nepl15* is transcriptionally active in embryonic (*Berkeley Drosophila Genome Project*), larval, and adult stages and enriched in metabolically relevant tissues, including the larval fat body (homologous to human liver and adipose tissue) and adult brain, gut, and heart, the organs central to nutrient sensing, storage, and systemic metabolic coordination (S. Banerjee et al., 2021). Furthermore, the single *Nepl15* transcript (Drucker & Banerjee, 2026) expression is higher in wild-type males than in females (S. Banerjee et al., 2021). Consistently, *Nepl15* knockout (*Nepl15^KO^*) whole adult male flies contained significantly reduced glycerolipids and glycogen reserves, while adult mutant females accumulated significantly more glycogen with a slight reduction in the lipids without affecting food intake as compared to the controls. The mutants displayed a delay in larval growth and a very high rate of mortality upon starvation. Additionally, the mutant adult males apparently exhibited reduced intracellular insulin signaling, but unchanged glucagon (Akh/ Adipokinetic hormone in flies) pathway (S. Banerjee et al., 2021). Glycogen and lipid metabolism, in coordination with the insulin/ *mTOR*, *Akh*, and *Sirtuin6* signaling pathways, overall controls fecundity, fertility, embryonic development, larval mortality, cellular and organismal growth, response to starvation and oxidative stress, climbing ability, heart function, aging, and lifespan of fruit flies, like in mammals. Altering the nutrient homeostasis in flies affects these traits, often in a sex-specific manner (S. Banerjee et al., 2021; Birse et al., 2010; Garrido et al., 2015; Heinrichsen & Haddad, 2012; Kannan & Fridell, 2013; Kanwal, Pillai, Samant, Gupta, & Gupta, 2019; Katewa & Kapahi, 2011; Y. Liu, Liao, Veenstra, & Nässel, 2016; Na et al., 2013; Roichman et al., 2021; Taylor et al., 2022; Teleman, 2009; Yamada, Habara, Kubo, & Nishimura, 2018a; Yamada et al., 2019). Thus, in the present study, we compared these characteristics between the *Nepl15^KO^* and isogenic *w^1118^* control flies in a sex and age-dependent manner.

We find that *Nepl15^KO^* flies display modest improvements in reproductive fitness, including laying a higher number of eggs, a higher percentage of pupariation, and a higher percentage of adult progeny eclosion. *Nepl15^KO^* adult female, but not male flies, exhibit extended lifespan when cultured on regular food. Both sexes show enhanced gut barrier integrity at 7 and 40 days of age, suggesting slower physiological aging in them. Age-related cardiac decline is attenuated, with preserved heart rate in older male and female mutants. Mutants show improved locomotor performance during progressive aging, with mutant females maintaining superior post-exercise performance and elevated ATP levels in the whole body and thorax. An indication of quick ATP turnover is observed only in 7-day-old mutant males as they exhibit a higher mitochondrial membrane potential. Consistently, the *Adenosine Monophosphate-activated Protein Kinase* (*AMPK*) shows reduced expression in both sexes of mutant flies. Furthermore, a significant decrease in total free radical production due to upregulation of *Superoxide dismutase 2* (*Sod2*) in the mutant females is likely to contribute to their cellular health and longer lifespan. Additionally, the marked reduction of *mechanistic Target of Rapamycin* (*mTOR*) expression and significant increase of *Sirtuin 6* (*Sirt6*) expression in mutant females are potentially contributing to lifespan extension cumulatively. In contrast, mutant males exhibit significantly reduced *mTOR* and *Sirt6* expressions, potentially limiting longevity benefits. Collectively, our results support a model in which *Nepl15* regulates nutrient reserves and metabolic signaling pathways, thereby linking energy homeostasis to lifespan, aging, activity, and cardiac and cellular health.

## Material and Methods

### Fly Strain and Fly Husbandry

Isogenic *Nepl15*-knockout (*w^1118^*; *Nepl15^KO^*) and *w^1118^* (as the control) flies were used in this study (S. Banerjee et al., 2021). All experiments were conducted using age-matched (5 - 7-day-old and 40 - 42-day-old) flies cultured under identical conditions, including keeping the population density constant. Stocks were maintained in an incubator (Percival Scientific DR-36NL) at 25 °C and 70% humidity on Nutri-Fly Pre-Mixed Fly Food (Genesee Scientific 66-113) supplemented with 20 mL 20% Tegosept (Methyl 4-hydroxybenzoate, Sigma H3647) and 20 mL propionic acid (Ward’s Science 470302-290) per 3.5L of food.

### Longevity assay

Newly eclosed mutant and control adult flies (n = 120) were sexed and housed at 10 male or female flies per vial on regular food. Flies were transferred to fresh food every 2–3 days, and deaths were recorded daily until all individuals died (Linford, Bilgir, Ro, & Pletcher, 2013). Survival data were analyzed using Kaplan–Meier survival analysis, and differences between survival curves were assessed using the log-rank (Mantel–Cox) test. This assay was performed in duplicate (> 200 flies per genotype per sex). Consistent results were observed across replicates; a representative set of data is shown.

### Smurf Assay

Intestinal barrier integrity was assessed using the Smurf assay on age-matched flies (7-day and 40-day old flies, n = 40 - 50 per genotype per sex) as previously described, with minor modifications (Martins, McCracken, Simons, Henriques, & Rera, 2018). One day before the assay, flies were transferred to food supplemented with either blue dye (Erioglaucine disodium salt, Sigma 861146) or Fluorescein sodium salt (Sigma F6377). Approximately 10 flies per genotype per sex were placed in each vial and allowed to feed at 25 °C for 16 hours. Following dye exposure, flies were anaesthetized with COLJ and imaged using a Leica EZ4W stereomicroscope. A fly was classified as a “Smurf” if dye was observed leaking outside the digestive tract into the abdominal cavity, whereas flies with dye restricted to the gut were classified as non-Smurfs. The percentage of Smurf-positive individuals was calculated for each sex and genotype.

### Optical coherence microscopy for heart function analysis

Adult flies were imaged using a custom-built SD-OCM system (Gracheva et al., 2022; Men et al., 2016; Men et al., 2020). Live flies were immobilized on the dorsal side up on the microscope slide with a small amount of rubber cement and placed on the mobile stage for imaging. The OCM light beam was directed at the A1 body segment. The images were generated with 128 A scans and 2000 B scans, an exposure time was set at 50 µs, and a frame rate was 125 frames per second, resulting in an approximately 16-second-long recording per sample. Raw OCM imaging data were processed using FlyNet3.0 software (Dong et al., 2020; Fishman et al., 2023; Ouyang et al., 2024).

### Climbing Assay

Climbing ability was measured using 10-, 20-, 30-, and 40-day-old adult flies. Groups of 10 flies per sex per genotype (total n = 100 flies per sex per genotype) were placed in an empty vial and gently tapped to the bottom. The number of flies that climbed 6 cm or above within 20 seconds was recorded (S. J. Banerjee et al., 2021). Each group underwent five trials, with one point awarded per successful climb per fly. The average climbing index was calculated as the total points divided by the number of flies used per trial and was used for statistical analysis.

### Exercise-induced climbing assay

Exercise-induced climbing ability was assessed using a rotating exercise paradigm (Watanabe & Riddle, 2018). Cohorts of adult flies (5-7 and 40-day-old; 10 flies per vial, per sex, per genotype) were transferred into clean, food-free conical tubes (Olympus Centrifuge Tubes 21-100) with 10 pinholes. Baseline locomotor performance was measured using the climbing assay as described above. Flies were then subjected to exercise in a 14-position rotating device (Tube revolver rotator, Thermo Scientific M5CT81001135) at 10 rpm for 45 min at 25 °C. Within 2 min of rotation cessation, climbing performance was reassessed under identical conditions. Seven biological replicates (n = 70 flies) were analyzed per sex per genotype, and the climbing index was calculated as described above.

### ATP determination

The measurement of ATP levels was performed using a luciferase-based assay kit (Molecular Probes ATP Kit; A22066) (Tennessen, Barry, Cox, & Thummel, 2014). ATP levels were used as a proxy for cellular energy status. Five adult flies (total 6 biological replicates, thus n = 30) or 15 thoraces (3 biological replicates) were collected and rinsed thoroughly with cold 1x phosphate buffer saline to remove food residues. The flies were homogenized in 100 µL Extraction buffer (6 M guanidine HCl, 100 mM Tris (pH 7.8), 4 mM EDTA), and 20 µL of the homogenate was set aside for protein quantification by Bradford Assay. The remaining homogenate was subjected to heat denaturation (Digital dry bath, Benchmark, Z742507) at 95 °C for 5 minutes, followed by centrifugation (Hermle ESCZ36-HK centrifuge, 12 x 1.5/2.0 mL rotor) at 13,000 rpm for 3 minutes at 4 °C to separate the supernatant. The supernatants were diluted 500-fold in dilution buffer (100 mM Tris and 4 mM EDTA, pH 7.8) and transferred to a white 96-well plate (Millipore, MSSWNFX40) for luminescence measurement. ATP standards were prepared from a 5 mM ATP stock solution to establish a standard curve for quantification according to the manufacturer’s instructions. Luminescence was measured immediately after the addition of 100 µL of luciferase reaction mix (8.9 mL dH_2_O, 0.5 mL 20x Reaction Buffer, 0.1 mL 0.1 M DTT, 0.5 mL of 10 mM D-luciferin, 2.5 μL of firefly luciferase 5 mg/mL stock solution for 100 reactions) to each well in a plate reader (Agilent BioTek Synergy H1). Three sequential measurements were averaged. The ATP concentrations in the samples were determined by comparing luminescence readings to the standard curve. The relative ATP levels were determined by normalizing luminescence values to total protein content, as measured using the Bradford assay.

### Mitochondrial membrane potential analysis

The mitochondrial isolation kit (Sigma, MITOISO1) was utilized for mitochondrial extraction and the JC-1 assay, following a previously established protocol (S. J. Banerjee et al., 2021). Mitochondria pellet prepared from 7-day-old, 40 adult *w^1118^* and *Nepl15^KO^*male and female flies per biological replicate (total of two biological replicates) was resuspended in 100 μL of ice-cold 1x Storage Buffer and maintained on ice. The isolated mitochondrial fraction was subsequently used for protein quantification by Bradford Assay. 20 μg of protein from each sample was used per 100 μL reaction of the JC-1 assay according to the manufacturer’s instructions to assess mitochondrial inner membrane potential.

### Quantification of total free radicals

Whole-body total free radicals were measured with the DCF ROS/RNS Assay Kit (Abcam ab238535) as described earlier (Arzoo, Tasmin, & Banerjee, 2025). For each biological replicate (total of three replicates), 5-7-day old, 25 adult control and mutant male and female flies were homogenized in 500 µL ice-cold 1x phosphate-buffered saline (10x PBS:1.37 M NaCl, 27 mM KCl, 100 mM Na_2_HPO_4_, 18 mM KH_2_PO_4_, pH 7.4). Homogenates were cleared by centrifugation at 10,000LJg (Hermle Z32-HK) for 5 min (4 °C); supernatants were split for protein and ROS/RNS determinations. Protein concentration was quantified using the Bradford assay (Bio-Rad 5000205). For the fluorescence assay, 50 µL of diluted lysate (1:4 dilution in 1x PBS) or hydrogen peroxide standard was pipetted in duplicate into a black 96-well plate (Thermo Scientific 165305), mixed with 50 µL 1x catalyst, and incubated for 5 min at room temperature. Subsequently, 100 µL DPS (containing DCF-DiOxyQ, Priming reagent and Stabilization solution) was added, followed by 45 minutes of incubation in the dark at room temperature. Fluorescence (ExLJ480LJnm, EmLJ530LJnm) was measured using an Agilent BioTek Synergy H1 plate reader. Raw relative fluorescence units were corrected by blank subtraction and normalized to soluble protein (FLULJmgLJ¹).

### Quantitative RT-PCR

Primers for qPCR were either designed using the Integrated DNA Technology qPCR PrimerQuest tool (Kalendar, 2025) or obtained from prior publications. They (see Supplementary Table 1) were tested for specificity and efficiency. To perform RT-qPCR, total RNA was extracted from tissues using TRI reagent (Sigma T9424). RNA was used for cDNA synthesis using iScript Reverse Transcriptase Supermix (Bio-Rad 1708841). RT-qPCR was performed using SsoAdvanced Universal SYBR Green Supermix (Bio-Rad 1725271) on a CFX Duet Real-Time PCR System (Bio-Rad). Target mRNA levels were normalized to the housekeeping gene *Ribosomal Protein L 32 (RpL32)*, and relative fold change of gene expression was determined using the 2^DΔΔCT^ method. All experiments were conducted in accordance with MIQE guidelines (S. Banerjee et al., 2021).

### Statistical Analysis

Sample sizes were determined based on previously published studies of *Drosophila* behavior and metabolism (S. Banerjee et al., 2021; Tennessen et al., 2014). All statistical analyses were performed using GraphPad Prism (version 9). Survival data were analyzed using the log-rank (Mantel-Cox) test (Fig. 2 a - b). For comparisons between (i) two groups, two-tailed unpaired t-tests (Fig. 1 i – k, 3 a – l, 4 a – h, 4 m – t, 5 a – d, 6 a – d, Supplementary Fig. 2B a – d, Supplementary Fig 3 a - d), and (ii) among multiple groups, one-way analysis of variance (ANOVA) (Fig. 3 m – n, 4 i - l), were performed. Data were presented as mean ± standard error of the mean (S.E.M) on bar graphs. Statistical significances were indicated as follows: \**P* < 0.05, ** *P* < 0.01, *** *P* < 0.001, and **** *P* < 0.0001; ns, not significant.

## Results

### 1. *Nepl15* deletion increases glycogen accumulation in ovaries and embryos, and modestly improves overall progeny productions

Glycogen and lipids deposited maternally into developing oocytes serve as essential energy reserves during embryogenesis and early larval development in *Drosophila* (Sieber, Thomsen, & Spradling, 2016; Song et al., 2019; Yamada et al., 2019). Alterations in embryonic glycogen or lipid stores can impair fecundity, fertility, embryonic development, reduce larval viability, and affect successful adult emergence (Tennessen & Thummel, 2011; Yamada, Habara, Kubo, & Nishimura, 2018b). Additionally, reduced fecundity and fertility were observed in flies storing excess TAG (triacylglycerol) (Na et al., 2013) or excess glycogen and lipid (Broughton et al., 2005). Because *Nepl15* is expressed in all embryonic stages of wildtype flies (Supplementary Fig. 1 a - d), and *Nepl15^KO^* females exhibited elevated whole-body glycogen levels (S. Banerjee et al., 2021), we first examined these nutrient reserves in ovaries and embryos, and next assessed the impact of these changes on reproductive performance, development, and survival of different life stages. Our data revealed that glycogen accumulation was visibly more in the *Nepl15^KO^* ovaries (Fig. 1 e, Supplementary Fig. 2B a) and embryos (Fig. 1 f - h, Supplementary Fig. 2B b) than in controls (Fig. 1 a, b – d, Supplementary Fig. 2B a, b), aligning with the mutant mother flies’ phenotype. However, the neutral lipid content did not show any changes in the ovaries (Supplementary Fig. 2A b, 2B c) and embryos (Supplementary Fig. 2A d, 2B d) of mutant female flies. Furthermore, from approximately 1,500 control eggs and 2,000 *Nepl15^KO^* eggs monitored throughout development, *Nepl15^K^* flies exhibited slightly higher median egg production (1091 vs. 922 eggs), pupation rate (59% vs. 54%), and adult eclosion rate (95.15% vs. 91.46%) than controls; however, these differences were not statistically significant (Mann–Whitney test; egg production, *P* = 0.6857; pupation, *P* = 0.6589; adult eclosion, *P* = 0.4586). Together, these findings indicate that loss of *Nepl15* increases glycogen accumulation in mutant females, ovaries, and embryos without adversely affecting reproductive performance or developmental progression.

**Figure 1.**
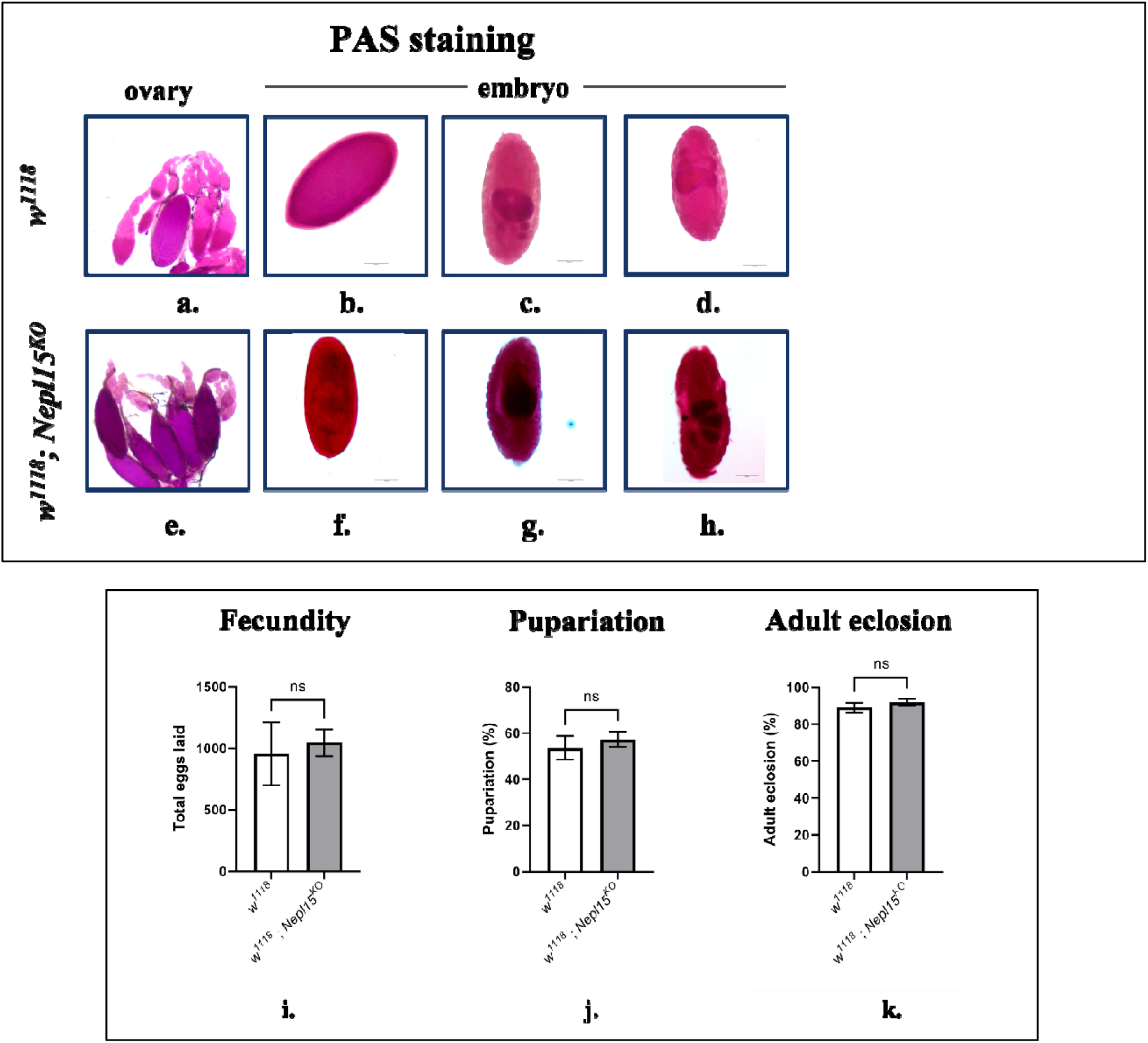
*Nepl15* deletion increases glycogen accumulation in ovaries and embryos and slightly increases reproductive fitness and pupal survival. Glycogen storage, as determined by PAS staining (pink), in the (a, e) ovaries (20 x) and embryos (20 x) of the (b - d) *w^1118^*control and (f – h) *Nepl15^K^*mutant flies (see supplementary method). (i) Fecundity, (j) percentage of pupariation, and (k) percentage of adult eclosion are all slightly higher in *Nepl15^KO^*flies (see supplementary method).

### 2. Loss of *Nepl15* prolongs the lifespan of female mutant flies and preserves intestinal barrier integrity of female and male mutant flies

Nutrient storage during adult life is a major determinant of organismal lifespan (Nasiri Moghadam et al., 2015). Genetic manipulation that reduced glycogen storage (Yamada et al., 2019) or enhanced glycogen and lipid reserves (Broughton et al., 2005) in adults, respectively reduced (Yamada et al., 2019) or expanded their lifespan (Broughton et al., 2005) substantially. Obese flies with high TAG storage showed drastically reduced lifespan (Chandegra, Tang, Chi, & Alic, 2017). We revealed that the deletion of *Nepl15* yielded a robust, female-specific extension of adult lifespan, but the longevity of mutant males was unaffected. In the first cohort, *Nepl15^KO^* females (n = 116) showed a median survival of 56 days, which was significantly more than the median survival of 39 days of *w^1118^* control female flies (n = 120) (Fig. 2 a). Another independent cohort (n = 107 *Nepl15^KO^*, 98 *w^1118^*) reproduced very similar results (*P* = 0.0003) (data not shown). In males (n = 119 *Nepl15^KO^*, 104 *w^1118^*) (Fig. 2 b), the mutant flies showed median survival of 47 days, which was more than that of the controls (45 days), but the overall difference in the lifespan was nonsignificant (Fig. 2 b). Similar results were observed in the other cohort of male flies (n = 117 *Nepl15^KO^*, 101 *w^1118^*) (*P* = 0.30) (data not shown). Thus, *Nepl15* ablation markedly prolong the female but not the male fly’s longevity when food is not scarce.

**Figure 2.**
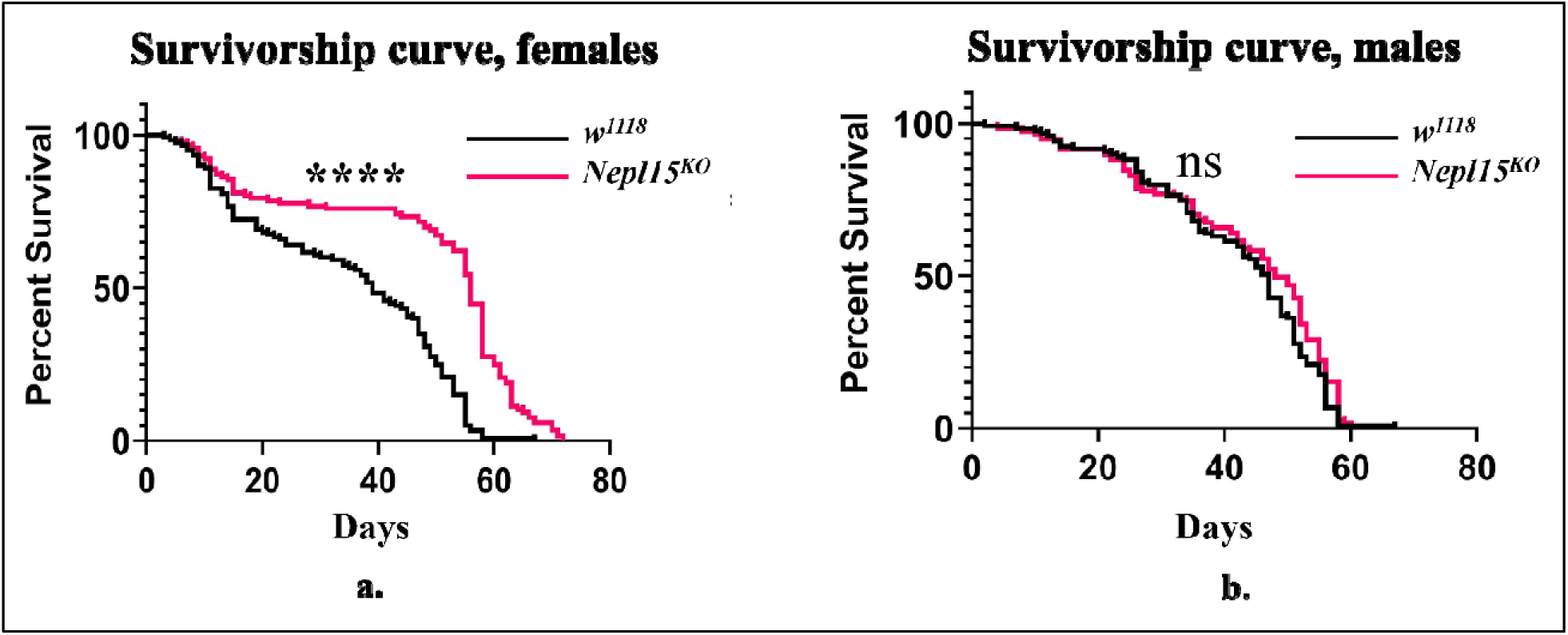
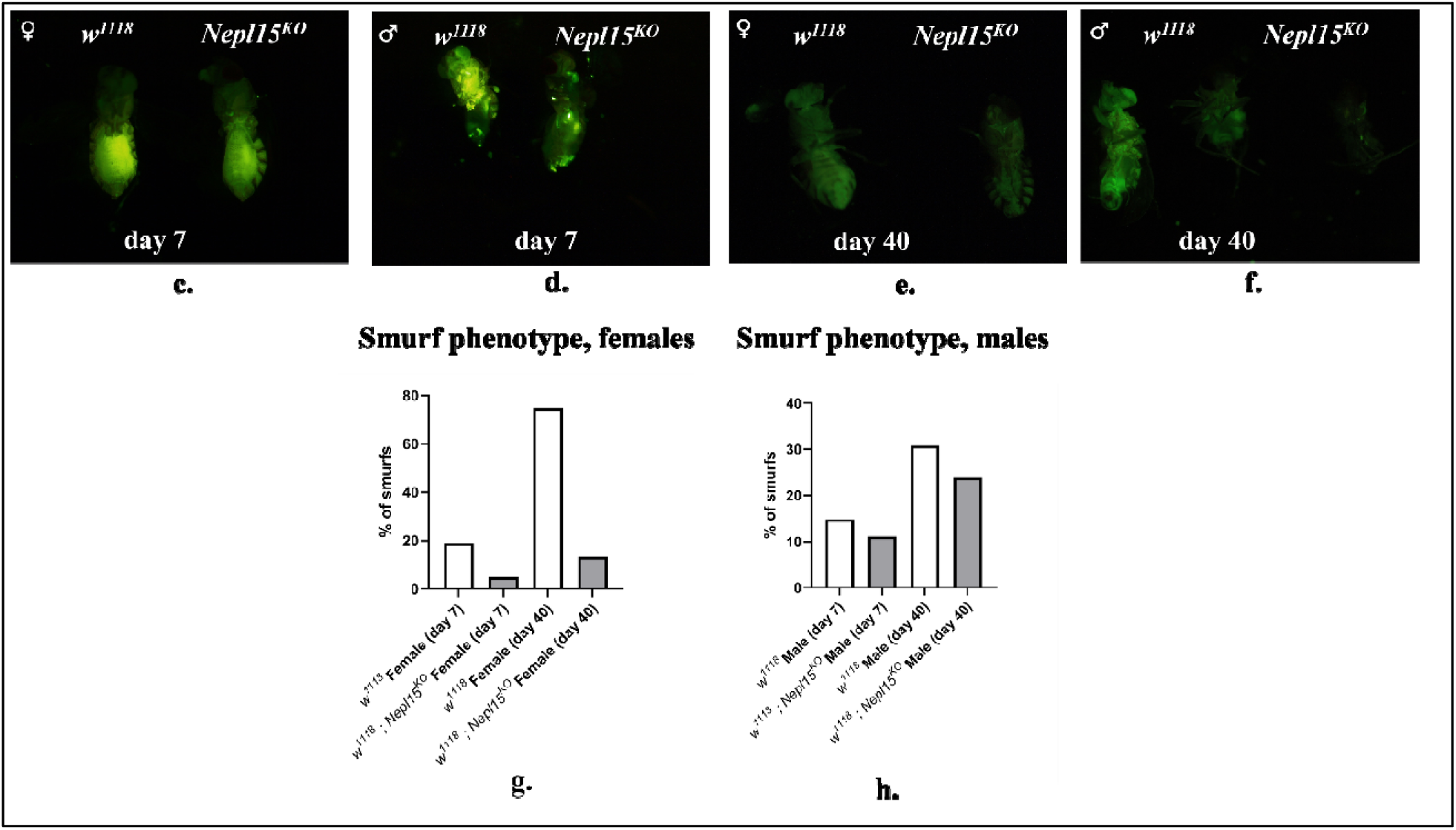
Knockout of *Nepl15* affects lifespan and gut barrier integrity. (a - b) Survivorship curves of *w^1118^* control (black) and *Nepl15^KO^*(magenta), female (a) and male (b) flies. (c - f) Representative image of control and *Nepl15^KO^* flies following ingestion of fluorescein-labelled food (green). (c) 7-day-old and (d) 40-day-old control and mutant females, and (g) corresponding female Smurf percentage. (e) 7-day-old and (f) 40-day-old control and mutant male flies and (h) corresponding male Smurf percentage.

Because *Nepl15* ablation extended female lifespan significantly, and male’s median lifespan slightly, we next examined whether mutants exhibited a relatively slower physiological aging. The gut is the primary nutrient-absorbing organ, and its barrier function is tightly linked to nutrient storage, metabolic stress, and late-life mortality (Salazar, Aparicio, Clark, Rera, & Walker, 2023). Reports indicate that aging or obesity-induced intestinal inflammation causes dysfunction of gut barrier integrity (Acciarino et al., 2024; Rera, Clark, & Walker, 2012). In flies, a major cause of death is age-dependent intestinal barrier failure, which allows gut microbes and toxins to enter the haemocoel and trigger systemic inflammation (Di Vincenzo, Del Gaudio, Petito, Lopetuso, & Scaldaferri, 2024). Thus, “Smurfness” (leaky gut) is widely used as a reliable marker for assessing the physiological aging of individual flies (Chelakkot, Ghim, & Ryu, 2018). Thus, we assessed gut permeability using the Smurf assay, where a non-absorbable dye ingested through food remains confined to the intestinal lumen unless epithelial integrity collapses. At 7-day age, 19% of *w^1118^* females showed Smurf phenotype, whereas less than 5 % *Nepl15^KO^* females showed Smurf (Fig. 2 g). By 40 days, the difference widened: 75% of control and 13 % of mutant females showed Smurf (Fig. 2 g). Male mutant flies also showed fewer Smurfs than controls, albeit to a lesser extent (Fig. 2 d, f). Smurf incidence rose from 14 % to 30 % in controls but only from 11 % to 24% in *Nepl15^KO^* male flies (Fig. 2 h) at 7 and 40 days of age, respectively. We repeated these experiments with non-fluorescent Brilliant Blue dye added to the food, and similar results were observed (the cumulative data were shown in Fig. 2 g and h). These data indicate that loss of *Nepl15* markedly delays age-associated intestinal barrier failure, particularly in females, indicating a healthier and slower physiological aging in the mutants.

### 3. *Nepl15* knockout flies exhibit better cardiac function with aging

Healthy aging is correlated with healthy heart function (Nishimura, Ocorr, Bodmer, & Cartry, 2011; Wessells, Fitzgerald, Cypser, Tatar, & Bodmer, 2004). Genetically mutated flies (Musselman & Kühnlein, 2018; Ocorr et al., 2007), as well as obesogenic flies with excess storage of glycerolipids due to intake of high-fat diets (Birse et al., 2010; Heinrichsen & Haddad, 2012) or high-sugar diets (Na et al., 2013) with enhanced insulin resistance (Chatterjee & Perrimon, 2021; Z. Liu & Huang, 2013), exhibited deteriorated heart functions (Birse et al., 2010; Heinrichsen & Haddad, 2012; Na et al., 2013). Given the improved gut barrier integrity and better lifespan observed in *Nepl15* knockout (*Nepl15^KO^*) flies (Smurf assay), we examined cardiac physiology in the *Nepl15^KO^,* which was likely to be altered due to altered nutrient storage-induced healthy aging. Moreover, this analysis was further motivated by high *Nepl15* expression in the wild-type flies, as shown in the FlyAtlas2 data (Leader, Krause, Pandit, Davies, & Dow, 2018). Cardiac parameters were quantified in control (*w^1118^*) and *Nepl15^KO^* flies at 7 and 40 days of age in both sexes, including heart rate (HR), end-diastolic area (EDA), end-systolic area (ESA), fractional shortening (FS), and arrhythmia index (AI). OCM of adult hearts revealed that removal of *Nepl15* reshapes heart geometry. At 7 days after eclosion, mutant males exhibited an increase in end-systolic area (ESA): from 3.3± 0.3 x 10^3^ µm^2^ in *w^1118^* to 4.5± 0.4 x 10^3^ µm^2^ in *Nepl15^KO^*, (*P* = 0.030), yielding a lower fractional shortening (FS) than the control (51.3 ± 3.1 % in *w^1118^* vs 40.5± 2.4 % in *Nepl15^KO^*, *P* = 0.006) (Fig. 3 b, d, f). The females showed the same trend: increased ESA, from 4.1± 0.7 x 10^3^ µm^2^ in *w^1118^* to 5.0± 0.9 x 10^3^ mm^2^ in *Nepl15^KO^* (not statistically significant, *P* =0.080), and reduced FS, 52.0 ± 2.1 % in *w^1118^* vs 45.4± 3.1 % in *Nepl15^KO^* (not statistically significant, *P* = 0.081) (Fig. 3 a, c, e). Aged flies (40 days after eclosion) EDA, ESA, and FS are summarized in Fig. 3 g - l. While young flies (day 7 after eclosion) of both sexes did not show the differences in EDA parameter (Fig. 3 a, b) between the two genotypes, we observed significantly smaller EDA in 40-day-old females (11.0± 0.6 x 10^3^ µm^2^ in *w^1118^*vs 9.2± 0.4 x 10^3^ µm^2^ in *Nepl15^KO^*, *P* = 0.024) and no differences in males (Fig. 3 g, h). ESA was similar in *w^1118^*and *Nepl15^KO^* males and females at day 40 after eclosion (Fig. 3 i, j). Ageing increased FS in both genotypes and sexes, indicating preserved compliance. However, FS in older *Nepl15^KO^* females was lower, 61.8 ± 1.4 % in *Nepl15^KO^* vs 66.7± 1.3 % in *w^1118^* (*P* = 0.014) (Fig. 3 k, l). Heart rate declines with age in control flies, whereas *Nepl15^KO^* flies, particularly mutant males, showed a profound preservation of heart rates between 7- and 40-day ages compared to controls. Arrhythmia index is significantly increased in aged (40-day-old) *Nepl15^KO^* females but remains unchanged in males at both ages, indicating a female-specific increase in cardiac rhythm instability with aging (Supplementary Fig. 3, Supplementary videos for heart data). Thus, the *Nepl15* mutant flies show resistance to age-dependent decline in cardiac functions, contributing to their healthy aging.

**Figure 3.**
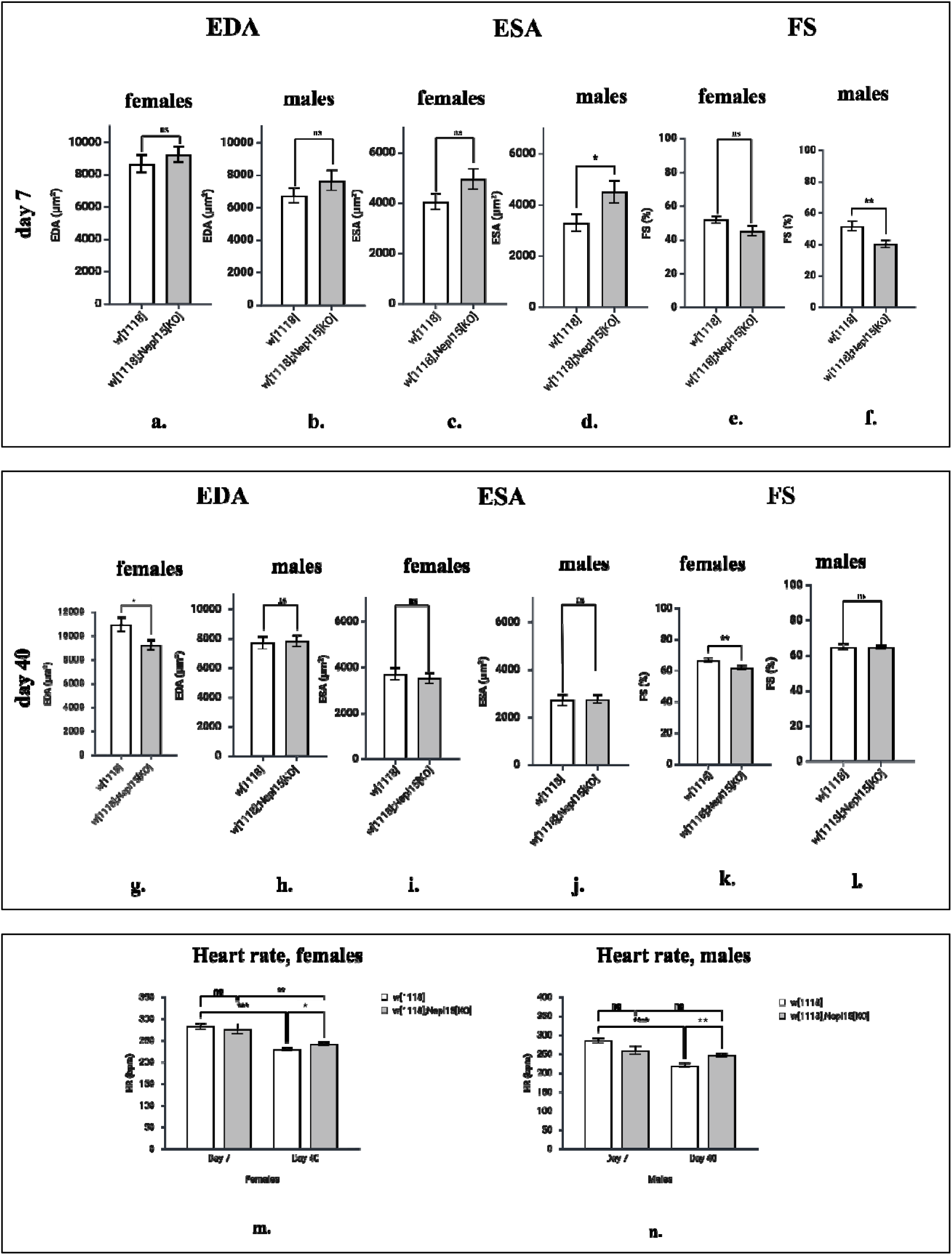
Effects of *Nepl15* deletion on cardiac contractile performance: OCM of adult *w^1118^*and *Nepl15^KO^* hearts were performed at day 7 (a - f) and day 40 (g - l) after eclosion. (a, b, g, h) End-diastolic area (EDA, μm²). (c, d, i, j) End-systolic area (ESA, μm²). (e, f, k, l) Fractional shortening (FS, %). (m, n) Heart rate (HR).

### 4. *Nepl15* knockout flies show higher and persistent climbing performance supported by sufficient energy availability

Whole-body ATP levels in the 7-day-old, whole body of control and mutant females (a) and males (b); and in the thorax of control and mutant females (m) and males (n). Mitochondrial membrane potential in the whole body of 7-day-old control and mutant females (q) and males (r). *AMPK*α expression in the whole body of 7-day-old control and mutant females (s) and males (t).

As the climbing ability declines with age, it has been used as a marker for aging in flies (Chatterjee & Perrimon, 2021; Gargano, Martin, Bhandari, & Grotewiel, 2005; Iliadi, Knight, & Boulianne, 2012; Sun et al., 2013). Moreover, this ability also declines in obesogenic flies with excess TAG storage (Birse et al., 2010), and in flies with reduced glycogen reserves (Yamada et al., 2019). Therefore, flies with poor nutrient storage or inefficient mitochondrial metabolism exhibit early declines in climbing capacity (Rhodenizer, Martin, Bhandari, Pletcher, & Grotewiel, 2008), whereas long-lived, nutritionally efficient genotypes maintain locomotor vigor into old age (Jones, 2012). Thus, we performed the negative-geotaxis climbing assay in an age- and sex-dependent manner, results of which could indicate anti-aging and anti-obesity benefits at the organismal levels in the *Nepl15* mutant flies. We monitored the climbing ability at four adult ages (days 10, 20, 30, and 40) in the control and mutant male and female flies (Fig. 4 a - h). At every time-point, the mean climbing index of female *Nepl15^KO^* cohorts exceeded that of *w^1118^*controls (Fig. 4 a - d), with an evidently increased performance at 40-days of age (Fig. 4 d; *P* = 0.019). (Fig. 4 e - h) The male *Nepl15^KO^* flies did not show any climbing decline but exhibited a non-significantly increased climbing ability at 40-day age only (Fig. 4 h; *P* = 0.6797). These findings indicate that *Nepl15* deletion sustains neuromuscular vigor deep into mid-life, avoiding the typical decline of climbing ability in the ageing control flies or other obese mutant flies (Birse et al., 2010).

**Figure 4.**
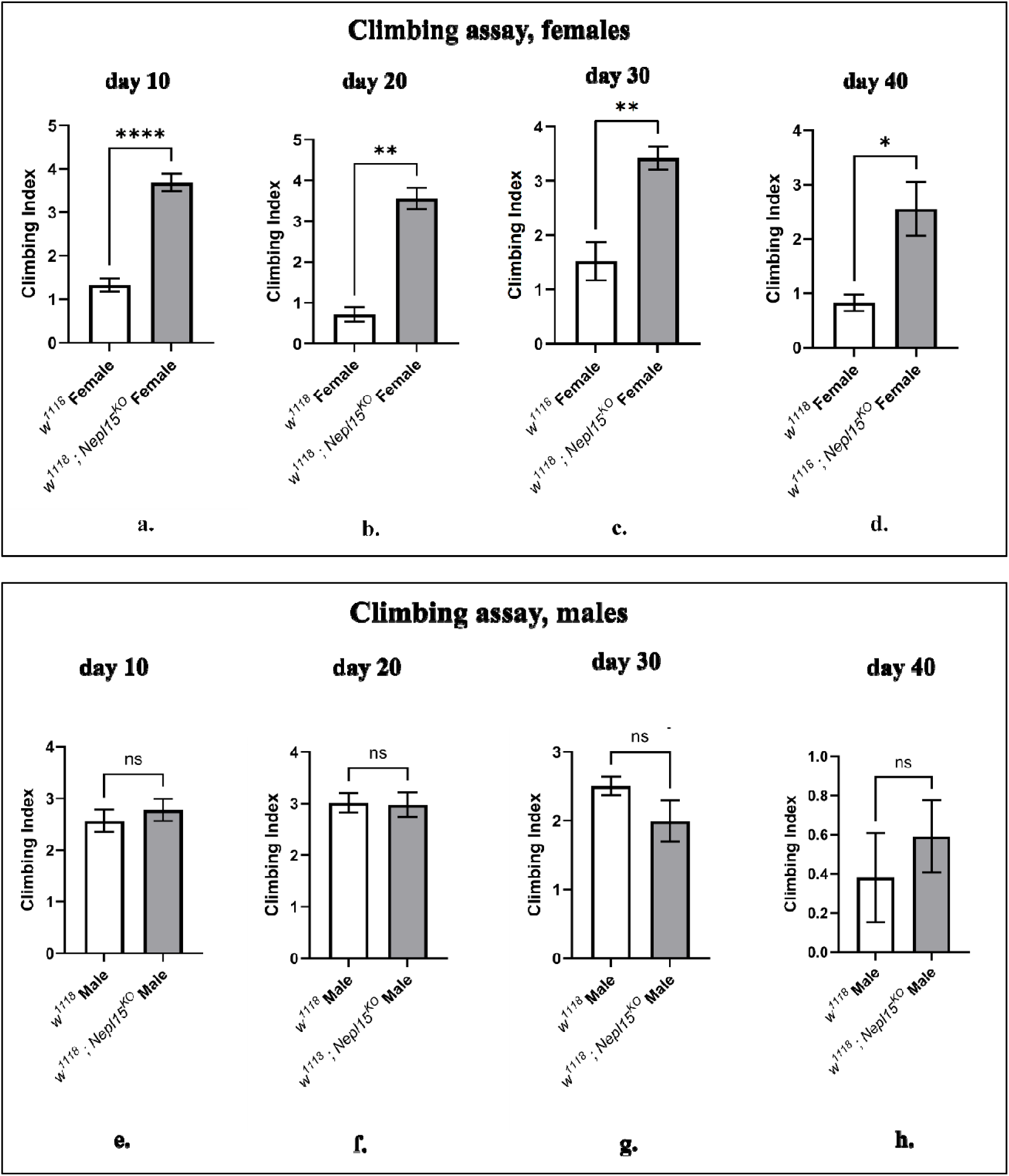
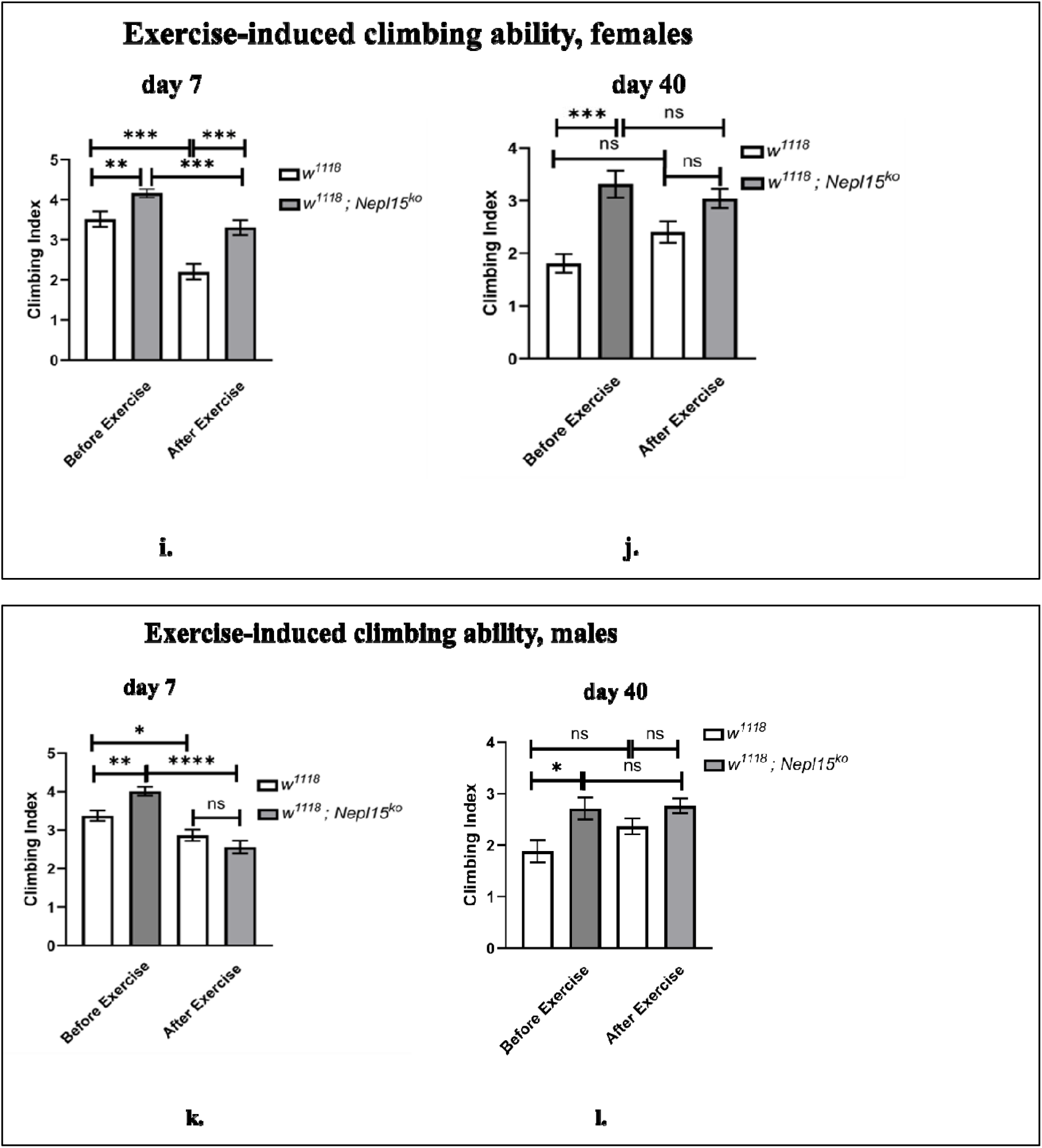
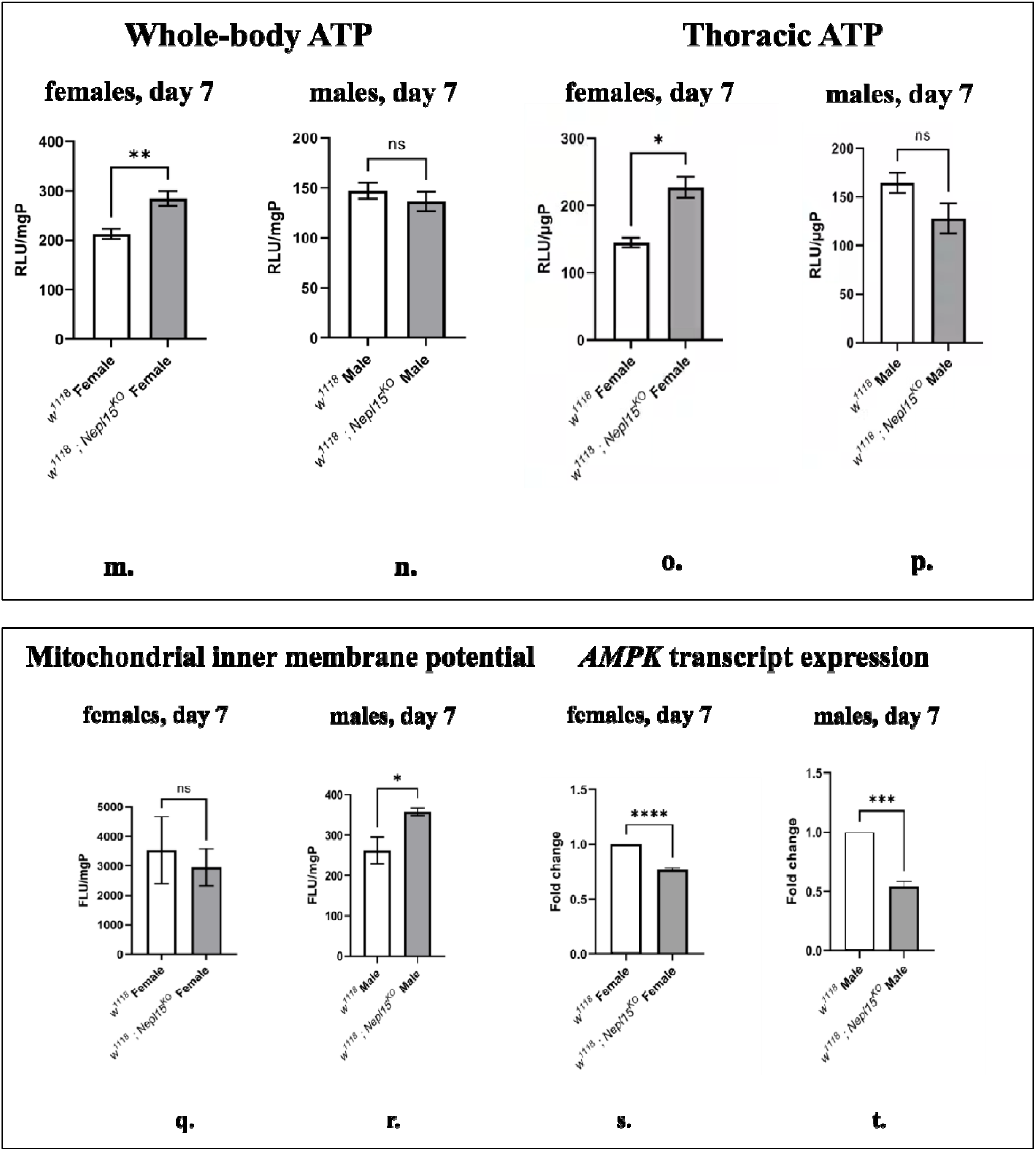
*Nepl15* knockout attenuates age-related decline in climbing ability and enhance postexercise climbing performance. (a - d) Climbing indices of female control and *Nepl15^KO^* flies at 10, 20, 30, and 40 days of age. (e - h) Climbing indices of male control and *Nepl15^KO^* flies at the corresponding ages. Climbing indices of (i) 5 to 7-day-old, and (j) 40-day-old female control and *Nepl15^KO^*flies, measured immediately before and after a 45-min rotational exercise regimen. Climbing indices of (k) 5 to 7-day-old, and (l) 40-day-old male control and *Nepl15^KO^* flies, measured immediately before and after a 45-min rotational exercise regimen. ***Nepl15* deletion changes energy homeostasis.**

We next examined whether this climbing ability persisted under acute energetic stress in *Nepl15^KO^* flies. We tested the climbing ability before and after challenging the animals with a 45-min exercise bout to test how efficiently they mobilize and utilize internal nutrient reserves, particularly relevant because *Nepl15* female mutants store less lipid yet more glycogen (S. Banerjee et al., 2021), a substrate that supports rapid ATP production during high-intensity activity (Winwood-Smith, White, & Franklin, 2020). (Fig. 4 i) The 5–7-day-old *Nepl15^KO^*female flies outperformed control flies’ climbing ability at pre-exercise (*P =* 0.0002) and post-exercise (*P =* 0.0008) periods, indicating quicker recovery and more efficient use of stored glycogen. (Fig. 4 j). At 40 days of age, the mutant females still showed significantly higher climbing ability (*P* = 0.0006) at the pre-exercise period, but there was no difference at the post-exercise period (*P* = 0.4720) compared to those of the age-matched controls, suggesting that their early-life metabolic resilience translates into preserved but no longer superior endurance in old age. Male climbing ability followed a different pattern. 7-day-old (Fig. 4 k) and 40-day-old (Fig. 4 l) *Nepl15^KO^* males displayed better pre-exercise climbing ability (*P* = 0.0016 and 0.0501) but, unlike females, did not show enhanced post-exercise climbing performances (*P* = 0.0743 and 0.5956). Their climbing index after exercise was indistinguishable from that of controls, consistent with reduced nutrient storage.

Locomotor activity in *Drosophila* depends heavily on ATP availability within muscle tissue (Chatterjee & Perrimon, 2021), and *Nepl15^KO^* flies maintained superior locomotor performances. ATP production depends on the efficient mobilization and utilization of stored nutrients (Chatterjee & Perrimon, 2021). The elevated glycogen storage observed in *Nepl15^KO^* females (S. Banerjee et al., 2021) might provide a rapidly accessible carbohydrate source during periods of high energetic demand, whereas reduced glycogen and lipid reserves in the mutant males might reduce ATP production. Thus, we investigated energy homeostasis in them. We quantified ATP levels in 7-day-old whole adult flies as well as in their isolated thoraces, which contain energy-demanding leg and flight muscles (Fabian, Schneeberg, & Beutel, 2016). Our results reveal significantly elevated ATP levels in *Nepl15^KO^* females’ whole body (Fig. 4 m) (*P* = 0.003) and thorax (Fig. 4 o) (*P* = 0.0413) relative to *w^1118^* controls at 7 days of age, aligning with excess glycogen reserves(S. Banerjee et al., 2021), and higher climbing ability in the mutant females compared to the controls. Surprisingly, the ATP levels in the whole body (Fig. 4 n) (*P* = 0.4271) and thorax (Fig. 4 p) (*P* = 0.1889) of the *Nepl15^KO^* male flies were not different from controls, even though the mutant males severely lack glycogen and lipid reserves (S. Banerjee et al., 2021). These findings indicate that loss of *Nepl15* does not impair energy availability but rather enhances ATP production in the mutant female flies.

ATP levels alone cannot distinguish how energy is produced (Zhang, Han, & Lin, 2018). Thus, we examined mitochondrial membrane potential (ΔΨm), a core determinant of oxidative phosphorylation efficiency. This potential reflects the proton-motive force that drives ATP synthesis; higher membrane potential typically indicates more efficient electron transport and a quicker ATP turnover, while a collapsed membrane potential signals mitochondrial dysfunction and disruption in ATP production (Ahmed Selim & Wojtovich, 2025). By measuring the mitochondrial membrane potential in isolated mitochondria, we could directly assess whether nutrient-storage alterations in *Nepl15* mutants translate into adaptive changes in mitochondrial energy-producing efficiency, providing a mechanistic bridge between altered metabolism and whole-body physiological performance. In 7-day-old adult flies, *Nepl15^KO^* males displayed a pronounced increase (Fig. 4 r) (*P* = 0.049) in the mitochondrial membrane potential, which was not seen in the mutant females (Fig. 4 q) (*P* = 0.784). This is likely the reason why the mutant male flies can produce comparable amounts of ATP, similar to the controls, to sustain their enforced climbing activity.

The AMP-activated protein kinase (AMPK) comprised of three subunits including the catalytic alpha (α) subunit, functions as a central cellular energy sensor whose activity is induced by a higher AMP/ATP ratio (Mihaylova & Shaw, 2011). We wanted to confirm that the cellular pathway in the mutants also sensed no decline in ATP levels in them by quantifying the *AMPK*α expression. Relative *AMPK*α transcript abundance was significantly reduced in the mutant female (Fig. 4 s) (*P* = 0.0001) and male (Fig. 4 t) (*P* = 0.0004) flies, compared to the corresponding 7-day-old control flies. Thus, cells of the mutant flies, especially the mutant males, are unlikely to experience energetic stress, and thereby do not trigger excess ATP production rate, which could further diminish their already low nutrient resources. Overall, these findings suggest that *Nepl15* deletion alters cellular energy homeostasis and nutrient-sensing pathways.

### 5. *Nepl15* loss reduces oxidative stress

In *Drosophila*, cellular oxidative stress exerted by the accumulation of free radicals (Reactive Oxygen Species/ ROS, and Reactive Nitrogen Species/ RNS) due to impaired antioxidant enzyme activity, aggravates cellular senescence (Orr, Radyuk, & Sohal, 2013), ultimately triggering age-associated physiological decline (Lennicke & Cochemé, 2020). Furthermore, diabetic patients with hyperglycemia show several pathological abnormalities, including cardiovascular and intestinal complications, that are triggered by ROS generation (Asmat, Abad, & Ismail, 2016; Fakhruddin, Alanazi, & Jackson, 2017; Giacco & Brownlee, 2010), indicating a direct relation between altered nutrient homeostasis and ROS-induced health decline. Thus, a longer lifespan is correlated with resistance to ROS (Piper & Partridge, 2018). Given that *Nepl15^KO^* flies showed multiple markers of healthier aging, we hypothesized that they possessed superior antioxidant mechanisms. Thus, we quantified the wholelJbody total freelJradical level and normalized it by protein amount in the same sample, and the expression levels of *Sod2*, the primary mitochondrial superoxide dismutase responsible for converting superoxide radicals generated during nutrient oxidation into less harmful species (Flynn & Melov, 2013; Islam, Alam, Islam, Nagar, & Rahman, 2026). At 7 days, *Nepl15^KO^*females exhibited a significant decrease in total free radicals (Fig. 5 a), whereas the mutant males showed a trend of decrease in total free radicals than the controls (Fig. 5 b). Thus, *Nepl15* deletion confers reduction in cellular oxidative stress during early adulthood, which is more prominent in the mutant females. Accordingly, we revealed that the loss of *Nepl15* elicited a sexlJbiased transcript expression of the *Sod2*. In 7-day-old mutant males, *Sod2* levels remain unchanged (*P* = 0.63; Fig. 5 c). By contrast, 7-day-old mutant females showed a robust ∼1.5-fold upregulation (*P* = 0.036; Fig. 5 d) of *Sod2* expression. These results aligned with the ROS and RNS levels observed in the mutant female and male flies.

**Figure 5.**
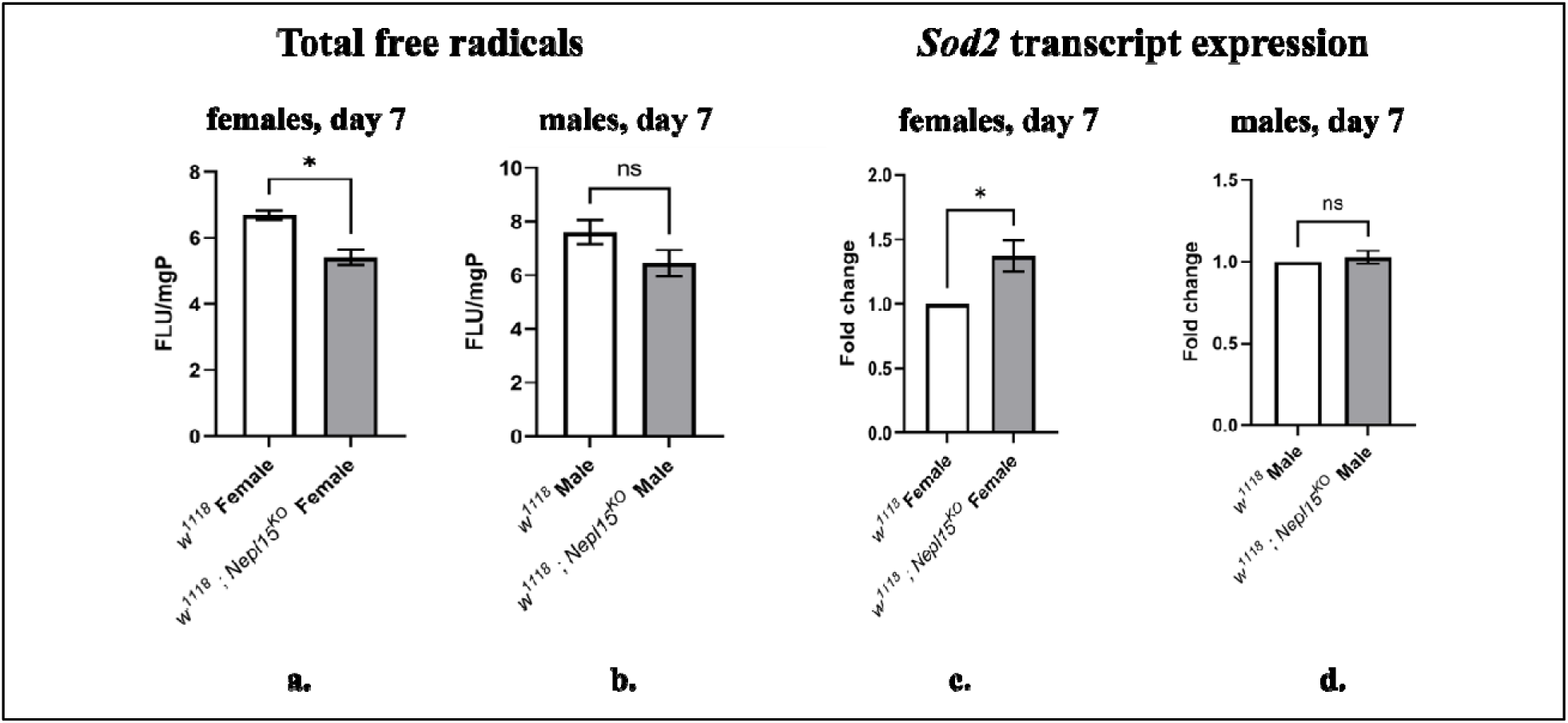
Lower free radical levels and *Sod2* expression in *Nepl15^KO^* female, but not in male flies. Total free radical levels in 7-day-old, whole adult control and mutant female (a) and male (b) flies. *Sod2* expression in 7-day-old, whole adult control and mutant females (c) and males (d).

### 6. *Nepl15* ablation suppresses□*mTOR* and alters *Sirt6* expressions

The *mTOR* gene encodes a conserved protein kinase that controls nutrient and energy metabolism, aging, and lifespan (Bjedov et al., 2010; Holczer et al., 2019; Piper & Partridge, 2018). Inhibition of mTOR function has been shown to extend lifespan in both mice and flies (Bjedov et al., 2010; Miller et al., 2014) and to delay cellular senescence through alleviating ROS production (Krishnamoorthy et al., 2018). Furthermore, cardiac dysfunction triggered by excess TAG reserves in obese flies can be rescued by inhibiting mTOR function (Birse et al., 2010). As the loss of *Nepl15* extended lifespan, preserved heart functions, and reduced ROS production, we examined the expression of *mTOR*. The *mTOR* transcript levels were significantly downregulated in 7-day-*Nepl15^KO^* female (Fig. 6 a) (*P* < 0.0001) and male (*P* = 0.0022) flies, aligning with our earlier observations on lifespan, ROS levels, and healthy aging in the mutant flies. However, the repression of *mTOR* expression could not extend the lifespan of the *Nepl15^K^*males, which aligns with the previous observations that suppression of mTOR activity increased the lifespan of female flies (Bjedov et al., 2010) and female mice (Harrison et al., 2009) more than the males, suggesting that additional regulatory pathways may contribute to the mutant fly lifespan. Thus, w examined *Sirt6* expression in these flies. *Sirt6* is a conserved NADlJ-dependent deacylase known to extend lifespan when overexpressed in both mice and *Drosophila* (Roichman et al., 2021), by promoting mitochondrial function, stress resistance, and genomic integrity (Taylor et al., 2022). In females, *Nepl15^KO^* mutants exhibited a significant increase in *Sirt6* expression compared to controls (Fig. 6 c) (*P* = 0.0147). Therefore, downregulation of *mTOR* and upregulation of *Sirt6* cumulatively extend the lifespan of mutant female flies. By contrast, male *Nepl15^KO^* flies showed a significant reduction in *Sirt6* expression relative to wild-type males (Fig. 6 d) (*P* = 0.002). Therefore, the simultaneous downregulation of both *mTOR* and *Sirt6* may offset each other’s effects on longevity in mutant males, ultimately preventing a significant extension of their lifespan.

**Figure 6.**
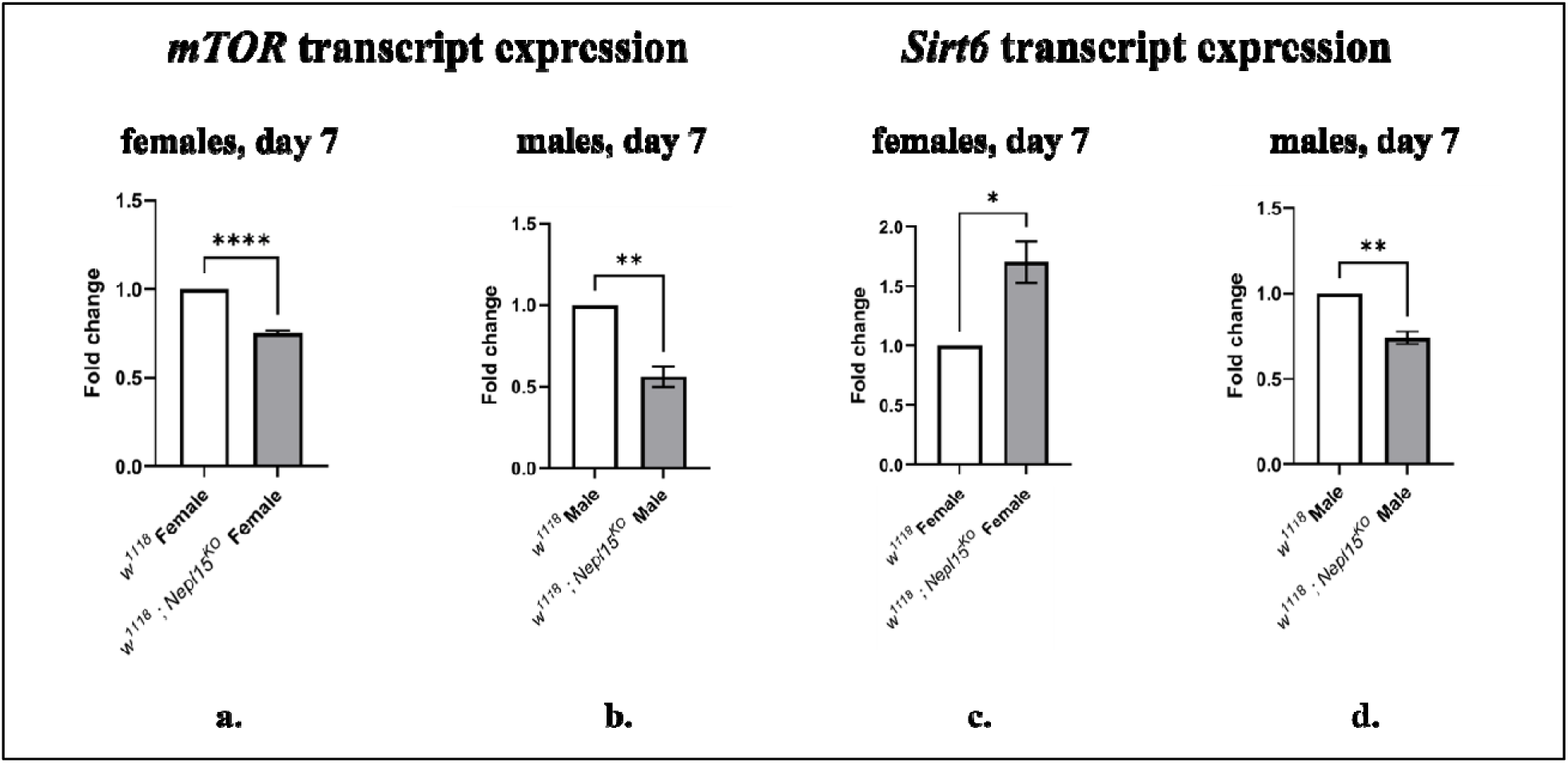
*Nepl15* deletion alters *mTOR* and *Sirt6* expressions in young adult flies. *mTOR* expression in 7-day-old, whole adult control and mutant females (a) and males (b). *Sirt6* expression in 7-day-old, whole adult control and mutant females (a) and males (b).

## Discussion

Genetic interventions altering glycogen storage in adult female flies similarly affected the maternally deposited glycogen levels in their embryos. Too high or too low glycogen content in these embryos caused substantial embryonic and larval deaths. Moreover, these embryos have altered metabolite profiles (Yamada et al., 2019). Additionally, adult flies with excess glycogen and lipid reserves had reduced fecundity (Broughton et al., 2005). Thus, changes in glycogen reserves in the adult female fly parent and their ovaries and embryos may produce detrimental effects on egg production, embryonic, and larval life. Interestingly, *Nepl15* gene deletion resulted in high glycogen reserves in adult mutant females, and extra glycogen allocation in their ovaries and embryos at such levels that did not reduce fecundity, fertility, larval, and pupal survival. We predict that *Nepl15* loss may also promote changes in other metabolites in the mutant embryos, which contribute to extra egg production, embryonic health, and survival.

In adult flies, less glycogen storage reduced lifespan; more glycogen and less lipid storage did not change lifespan (Yamada et al., 2019); excess glycogen and lipid storage increased lifespan (Broughton et al., 2005); and excess lipid storage plus more circulating carbohydrate reduced lifespan (Chandegra et al., 2017). Thus, changes in the nutrient levels alone cannot explain their effects on lifespan. Genetic interventions or feeding behavior that triggered these changes, as well as the relative levels of different nutrients and other physiological factors, are likely to contribute to lifespan. *Nepl15^KO^* female flies exhibited significantly longer lifespan, while the mutant males also showed a slightly better median lifespan. Therefore, we envisioned that additional cellular, organ, and organismal level benefits are likely to promote the lifespan of *Nepl15* mutant flies. Accordingly, lower ROS levels, higher ATP levels or better mitochondrial membrane potential that can trigger a quick ATP turnover, better gut barrier integrity, preserved heart function, enhanced locomotor ability with progressive age, all are likely to contribute positively to overall health and lifespan of the mutant flies.

Consistent with these health benefits, at the cellular signaling level, *Nepl15^KO^* flies showed reduced *mTOR* expression, which is known to extend lifespan (Bjedov et al., 2010; Miller et al., 2014) and to ameliorate cardiac arrhythmia resulting from lipid accumulation due to high-fat diet intake by flies (Birse et al., 2010). However, reduced mTOR function has also been reported to reduce climbing ability (Birse et al., 2010), and male and female fertility (Oliveira, Cheng, & Alves, 2017), which were not true for the *Nepl15* mutant flies, suggesting that *Nepl15* ablation might alter the pathways triggered by the inactivation of mTOR which otherwise impair the locomotor ability and fertility. Additionally, overexpression of *Sirt6* in the *Nepl15* mutant adult female flies was also likely to enhance their lifespan (Taylor et al., 2022). Intriguingly, the downregulation of *Sirt6* in the *Nepl15* mutant adult male flies was unable to reduce their lifespan, which requires further investigations to understand the detailed mechanism.

Overall, this study demonstrates that, consistent with the sex-biased expression of *Nepl15* in wild-type (FlyAtlas2) and *w^1118^* flies, and with the sex-specific alterations in nutrient and energy storage caused by *Nepl15* deletion in adult mutant females and male flies (S. Banerjee et al., 2021), *Nepl15* mutants also exhibit several sexually dimorphic cellular and physiological phenotypes. These phenotypes strongly indicate that anti-aging and anti-obesity processes are promoted due to the deletion of the *Nepl15* gene, which could have substantial implications for maintaining a healthy life in the future.

## Supporting information

Supplementary result and methods

Supplementary videos for heart data

## Author contribution

S.H.A.: Conceptualization, experimental design, methodology, investigation, validation, formal analysis, data curation, visualization, writing, original draft preparation, review and editing.

C.D.: Methodology, experimental design, data curation, visualization, writing, investigation, formal analysis, review and editing

N.J.: Investigation and formal analysis.

S.J.B.: Conceptualization, experimental design, supervision, project administration, resources, methodology, investigation, validation, formal analysis, data curation, visualization, writing, original draft preparation, review and editing, and funding acquisition.

E.G., A.M., R.H., F.W., and C.Z.: Cardiac physiology experiments and analysis, including heart imaging and quantitative assessment.

All authors have read and agreed to the published version of the manuscript.

## Acknowledgement

All fly husbandry, molecular experiments, and imaging (excluding cardiac physiology assays) were carried out in the Department of Biological Sciences, Texas Tech University. Cardiac physiology experiments were performed at Washington University in St. Louis.

## Data Availability Statement

The data that support the findings of this study are available from the corresponding author upon reasonable request.

