## Supplementary result and methods for "*Drosophila melanogaster Nepl15* gene deletion promotes anti-aging and anti-obesity phenotypes in a sex-dependent manner"

**Supplementary Data**

**1. *Nepl15* mRNA is ubiquitously expressed in all embryonic stages of wild-type flies**

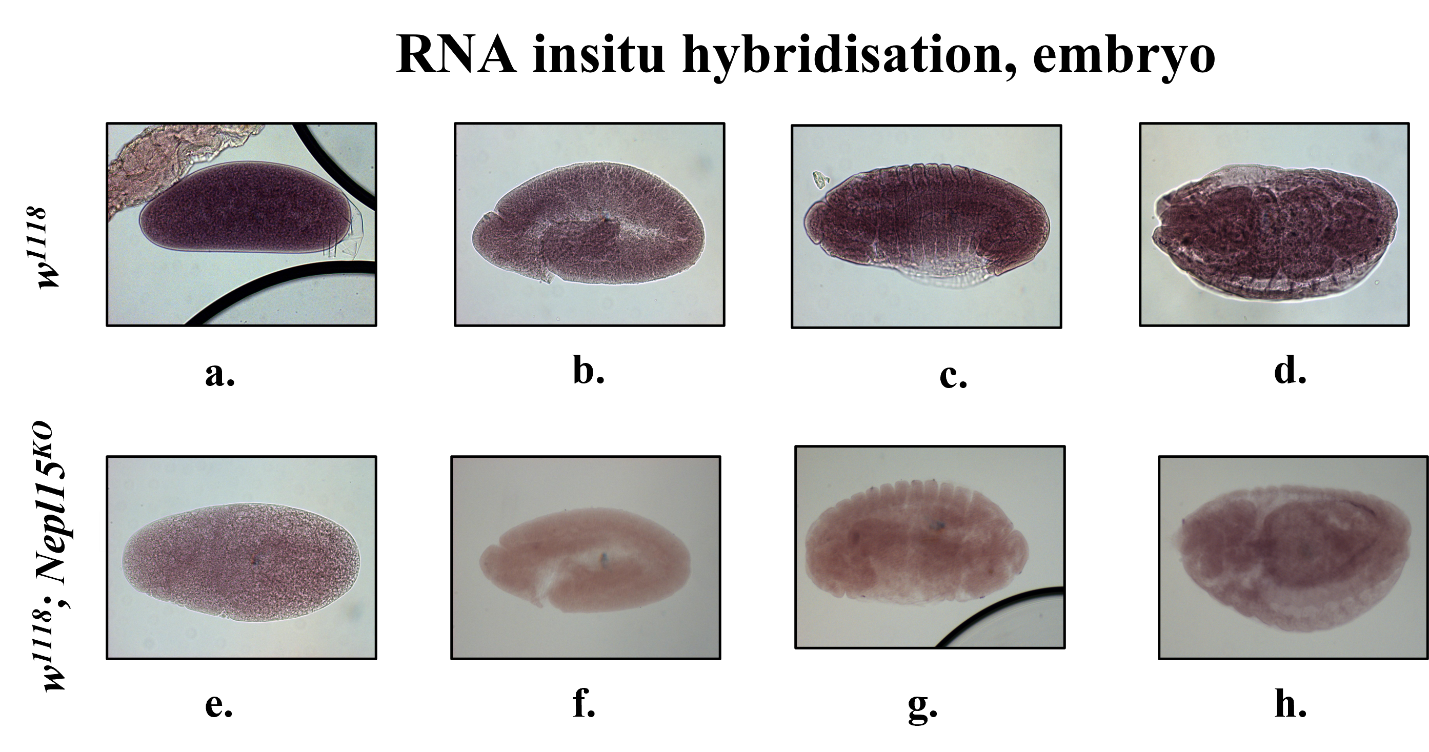

**Supplementary Figure 1.** ***Nepl15* mRNA expression is observed in all embryonic stages of wild-type flies.** Digoxigenin-labeled specific antisense probes marking the (a-d) presence of *Nepl15* mRNA (dark brown) in wildtype embryos, and (e-h) absence of *Nepl15* mRNA (light brown) in *Nepl15^KO^* embryos by in situ hybridization.

**2. *Nepl15^KO^* does not change the lipid reserves in the ovaries and embryos**

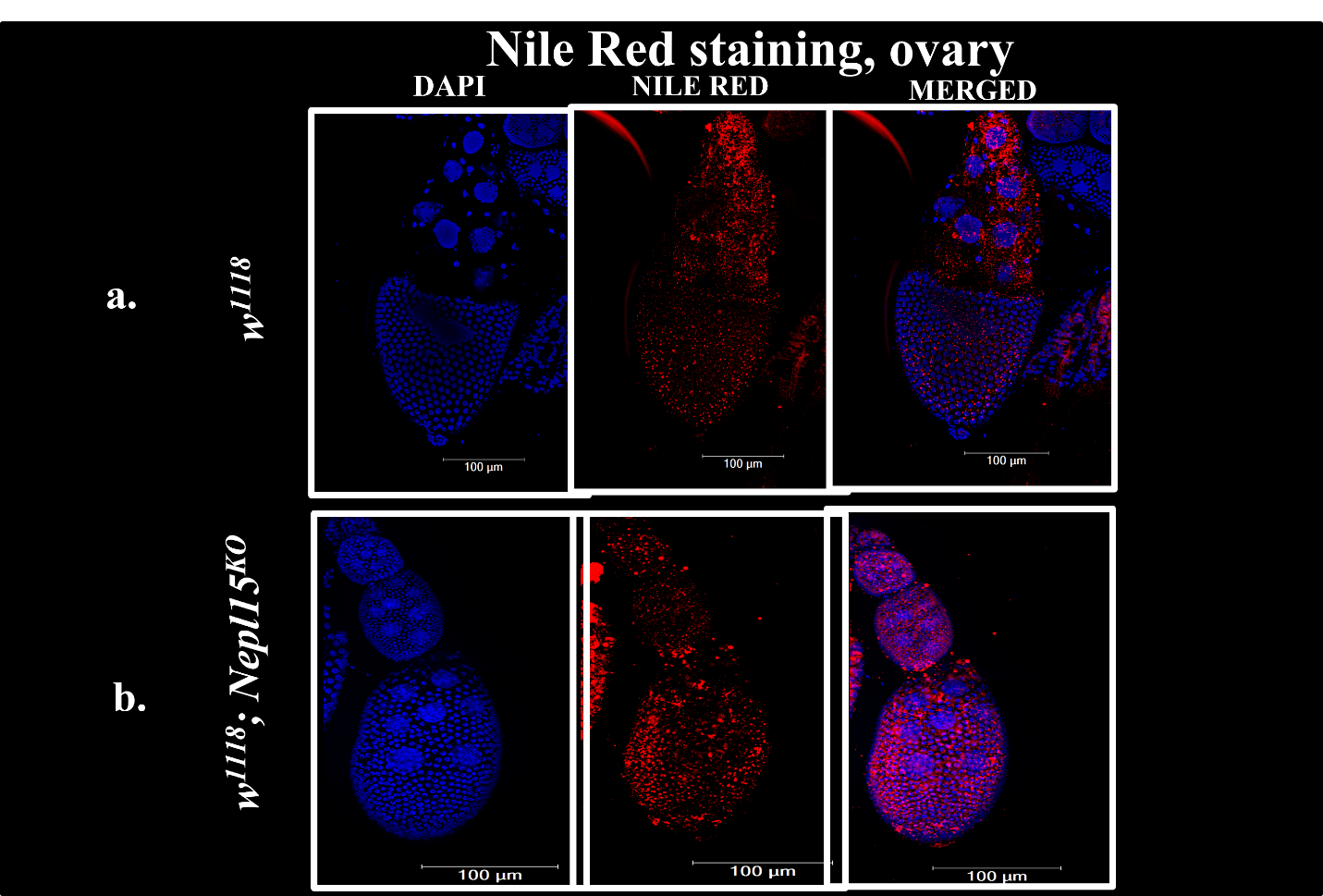

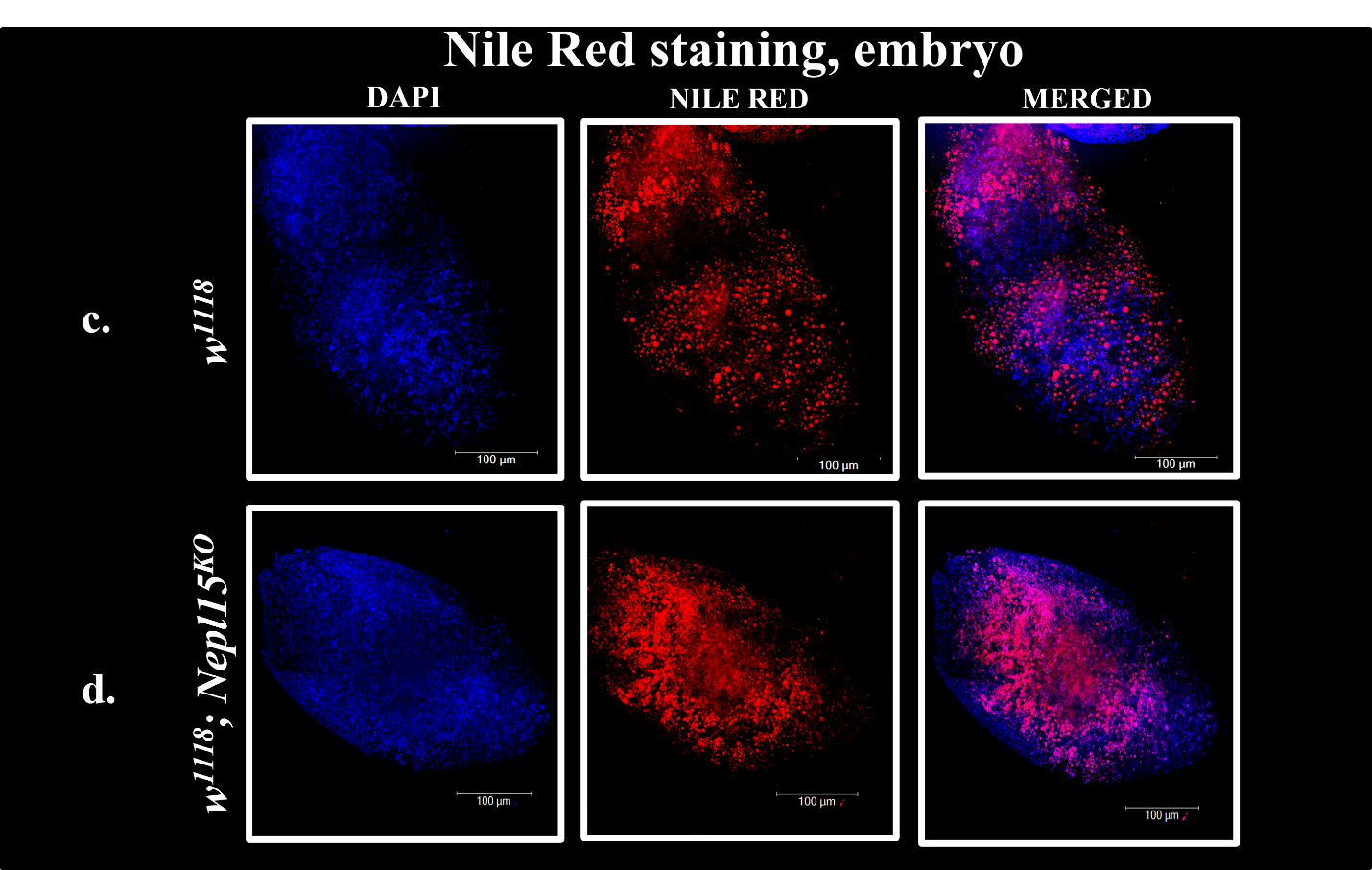

**Supplementary Figure 2A.** **No changes were observed in the lipid reserves in the *Nepl15^KO^* ovaries and embryos.** Nucleus (Dapi staining, blue), Neutral lipid in lipid droplets (Nile Red staining, red), and merged images of ovaries (Fig. 2A a, b) and embryos (Fig. 2A c, d) of control and mutant flies.

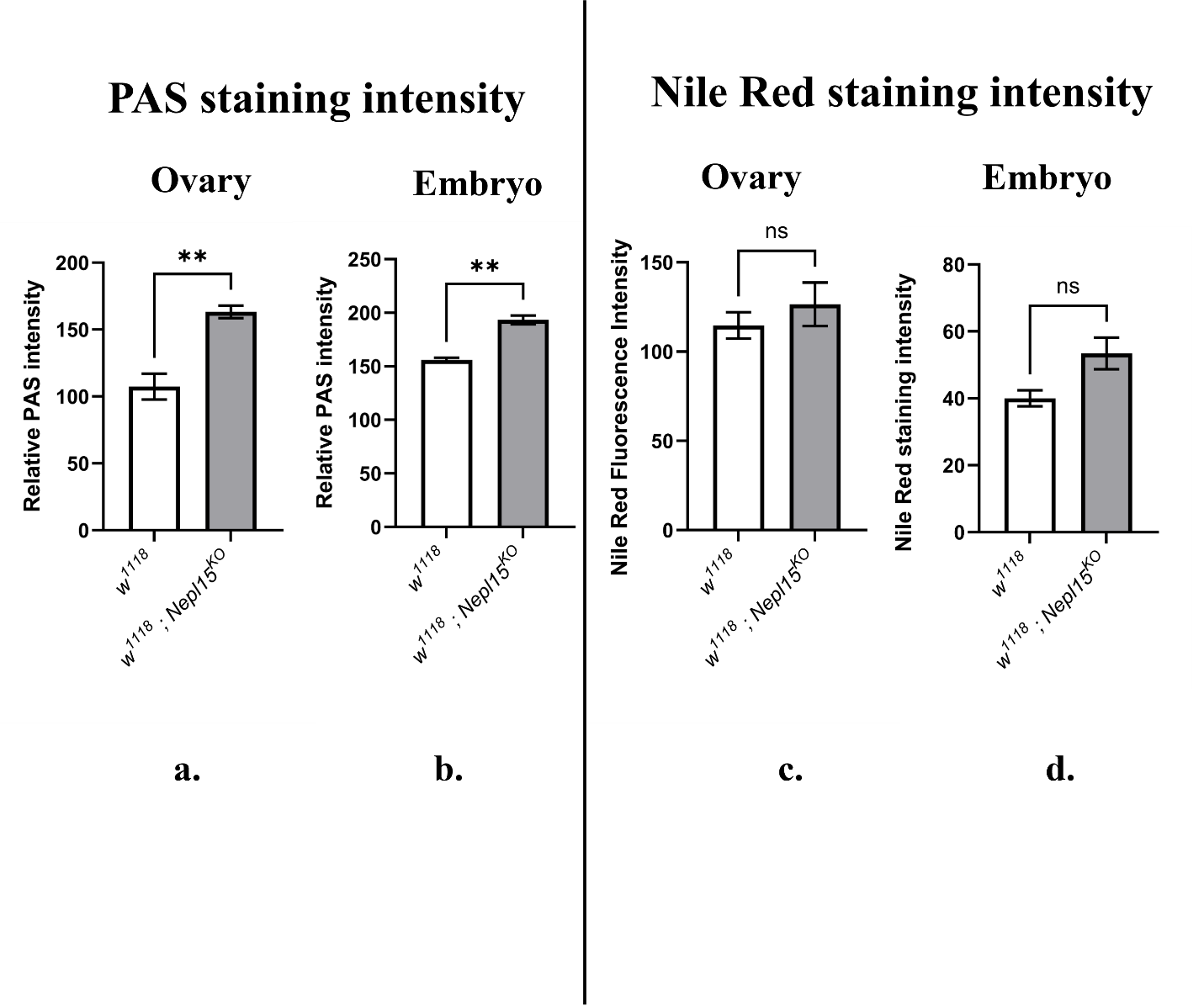

**Supplementary Figure 2B. Quantification of PAS (marks glycogen reserves) signal and Nile Red (marks neutral lipids in lipid droplets) signal intensities in embryos and ovaries of *w^1118^* control, and *Nepl15^KO^* mutant flies.** Relative PAS signal intensity was quantified in ovaries (a), embryos across all stages from (b) control (*w¹¹¹⁸*) and *Nepl15^KO^* flies. Relative Nile Red signal intensity was quantified in ovaries (c), embryos across all stages from (d) control (*w¹¹¹⁸*) and *Nepl15^KO^* flies.

**3. Loss of *Nepl15* increases the arrhythmia index only in 40-day-old mutant females**

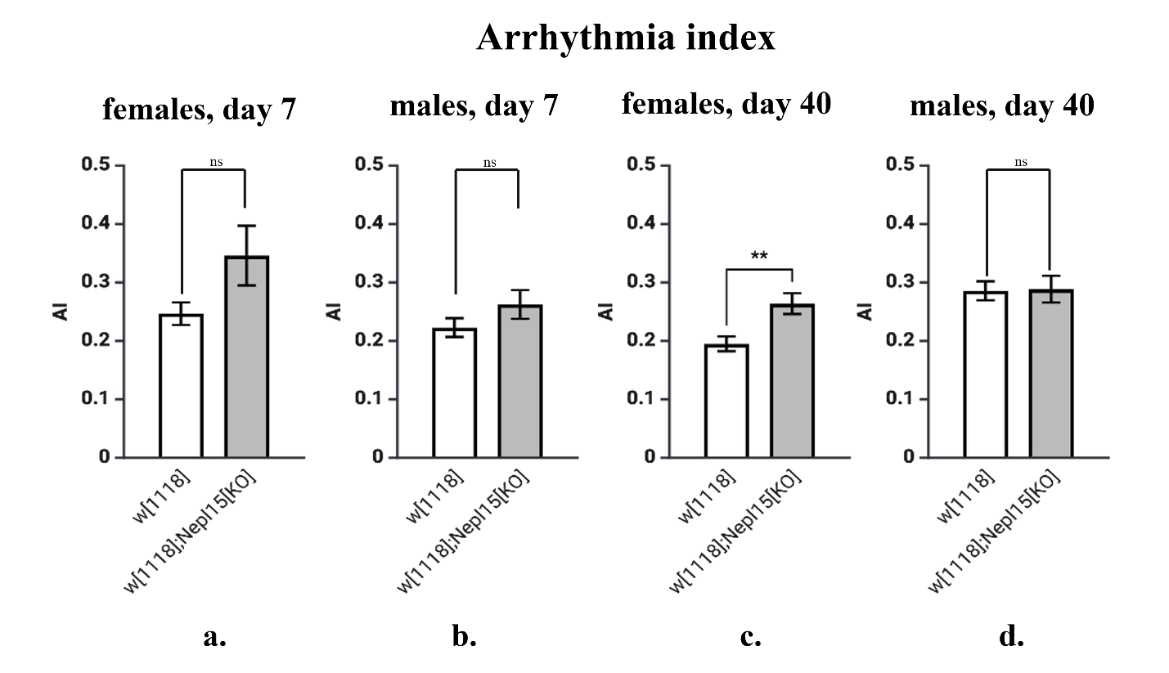

**Supplementary Figure 3. Effects of *Nepl15* deletion on cardiac arrhythmia index (AI) during aging.** Arrhythmia index (AI) was quantified in *w^1118^* and *Nepl15^KO^* flies at 7 and 40 days of age in both sexes using optical coherence microscopy-based heart recordings. AI in 7-day-old (a) female, and (b) male; and 40-day-old (c) female and (d) male control and *Nepl15^KO^* flies.

**Supplementary Methods**

**RNA in situ hybridization for detecting *Nepl15* mRNA at embryonic stages.** Digoxigenin-labelled (Roche 11175025910) sense and antisense probes for RNA in situ hybridization were prepared and performed following a published protocol (Firth & Baker, 2007). All stages of embryos collected from *w^1118^* (positive control) and *Nepl15^KO^* mutant (negative control) flies were dechorionated in 50% bleach, and fixed in heptane/ 4% paraformaldehyde, followed by methanol treatment for devitalization. Embryos were pre-hybridized and hybridized overnight at 55°C with denatured sense and antisense probes (1:1000), washed, and incubated with alkaline phosphatase–conjugated anti-DIG antibody. Signals were developed using NBT/BCIP. Embryos were mounted in 80% glycerol and imaged by bright-field microscopy. No signal was found in the samples stained with the sense probes. Antisense probes produced strong signals (dark brown color) only in the *w^1118^* embryos, but not in the mutant embryos (weak signal, light brown color).

**Histological analysis of glycogen and lipid storage in embryos and ovaries.**

Ovaries and embryos were collected for histological analysis of glycogen and neutral lipid storage. Ovaries were dissected in ice-cold 1× phosphate-buffered saline (PBS), whereas embryos were collected on egg-lay plates, dechorionated in 50% bleach for 2 min, washed thoroughly with water, and devitellinized using a heptane-methanol procedure (Kilwein, Dao, & Welte, 2023). Embryos and ovaries were fixed in freshly prepared 4% paraformaldehyde (PFA) in 1× PBS for 20–30 min at room temperature and subsequently washed three times with ice-cold 1× PBS.

For glycogen visualization, fixed samples were incubated in 1% periodic acid solution (Sigma-Aldrich 3951) for 5 min at room temperature, followed by three washes with PBS containing 1% bovine serum albumin (BSA). Samples were then immersed in Schiff’s reagent (Sigma-Aldrich 3952) for 15 min at room temperature and washed three additional times with PBS containing 1% BSA. PAS-positive staining was used to visualize glycogen and other polysaccharide-containing structures. Samples were mounted in 80% glycerol and imaged using bright-field microscopy (Banerjee et al., 2021).

For neutral lipid staining, separate fixed samples were incubated for 30 min at room temperature in a staining solution containing Nile Red and Hoechst 33342. Nile Red stock solution (0.05%) was prepared by dissolving 5 mg Nile Red (Sigma-Aldrich 72485) in 10 mL methanol and stored protected from light at 4°C. Nile Red was used at a final dilution of 1:2000 in 1× PBS. Hoechst 33342 (Sigma-Aldrich 14533) working solution was prepared by diluting the intermediate stock 1:10 in PBS and subsequently used at a final dilution of 1:100. Samples were protected from light throughout the staining procedure by wrapping tubes in aluminum foil. Following staining, tissues were washed three times with ice-cold 1x PBS and mounted in 80% glycerol containing N-propyl gallate as an antifade reagent (Banerjee et al., 2021).

PAS-stained samples were visualized using bright-field microscopy, whereas Nile Red-stained samples were imaged using a Leica DMi8 Inverted THUNDER Confocal Spinning Disk microscope. Nile Red fluorescence was detected using an excitation wavelength of 485 nm and an emission wavelength of 595 nm. Hoechst 33342 fluorescence was detected using an excitation wavelength of 361 nm and an emission wavelength of 486 nm. For each genotype, Z-stack images were acquired from at least five biological samples using identical imaging settings across the experimental group (Banerjee et al., 2021).

**Image analysis for quantification of PAS and Nile Red signal intensities**

PAS- and Nile Red-stained samples were imaged under identical microscope settings for each experiment. Image analysis was performed using Fiji/ImageJ (National Institutes of Health, Bethesda, MD, USA). For z-stack images, maximum-intensity projections were generated from all optical sections before analysis. For Nile Red fluorescence images, the appropriate fluorescence channel was isolated by splitting the image channels.

A rectangular region of interest (ROI) of identical size was placed over the corresponding anatomical region in each embryo or ovary. The same ROI was applied to all images within an experiment to ensure consistency. Mean pixel intensity was obtained from the histogram function in Fiji/ImageJ and used for quantitative analysis. Images from control and mutant samples were processed using identical acquisition parameters and image analysis settings. Mean intensity values from independent biological samples were exported to Microsoft Excel for statistical analysis (Banerjee et al., 2021).

**Fecundity, Pupariation, and Adult Eclosion Assays**

Groups of freshly emerged *w¹¹¹⁸* and *Nepl15^KO^* flies, each consisting of 20 females and 10 males, were aged for four days in vials containing standard cornmeal food at 25 °C (Banerjee et al., 2021). Flies were then transferred to four egg-laying chambers per genotype containing egg-laying medium [4% (w/v) *Drosophila* agar (USBiological Life Sciences A0940) and 40% (v/v) molasses (Sweet Harvest Foods Golden Molasses Unsulphured)] supplemented with freshly prepared yeast paste (Fleischmann's Instant Dry yeast mixed with water to a toothpaste-like consistency) and allowed to oviposit for four consecutive days. Eggs laid by 5–7-day-old flies were collected daily, and approximately 100 randomly selected eggs were seeded into each vial containing standard food. In total, approximately **1,500** *w¹¹¹⁸* **eggs** and **2,000** *Nepl15^KO^* **eggs** were monitored throughout development. The numbers of pupae and subsequently eclosed adults were recorded to determine egg-to-pupa survival (pupation rate) and pupa-to-adult survival (adult eclosion rate). Fecundity was assessed by manually counting the total number of eggs laid by each genotype over the four-day collection period under a stereomicroscope (Leica EZ4 W).

**Supplementary Table 1: Forward (FP) and reverse (RP) primer sequences used for RT-qPCR.**

| Target genes | Sequence of Primers |
| --- | --- |
| *RpL32* (Banerjee et al., 2021; Joshi, Banerjee, Curtiss, & Ashley, 2022) | FP-CCAAGCACTTCATCCGCCACC  RP-GCGGGTGCGCTTGTTCGATCC |
| *Sod2* | FP-AGAGCCTCGAGGAGTTCAAA  RP-CAGTTTGCCCGACTTCTTGT |
| *mTOR* | FP-CGCAGGCTCTGGTCTATC  RP-AGGTGGGCGAGTGTTTC |
| *Sirt6*  (Mishra & Mishra, 2024) | FP-GCGGATGGATTG TCAGCCTA  RP- GAGGACAACGTGTCCCGATT |
| *AMPK**α* | FP- GACCGTTGGCGGAGGTA  RP- CGGATGCGGTCGAATGG |
