## Supplementary videos for heart data for "*Drosophila melanogaster Nepl15* gene deletion promotes anti-aging and anti-obesity phenotypes in a sex-dependent manner"

### Slide 1
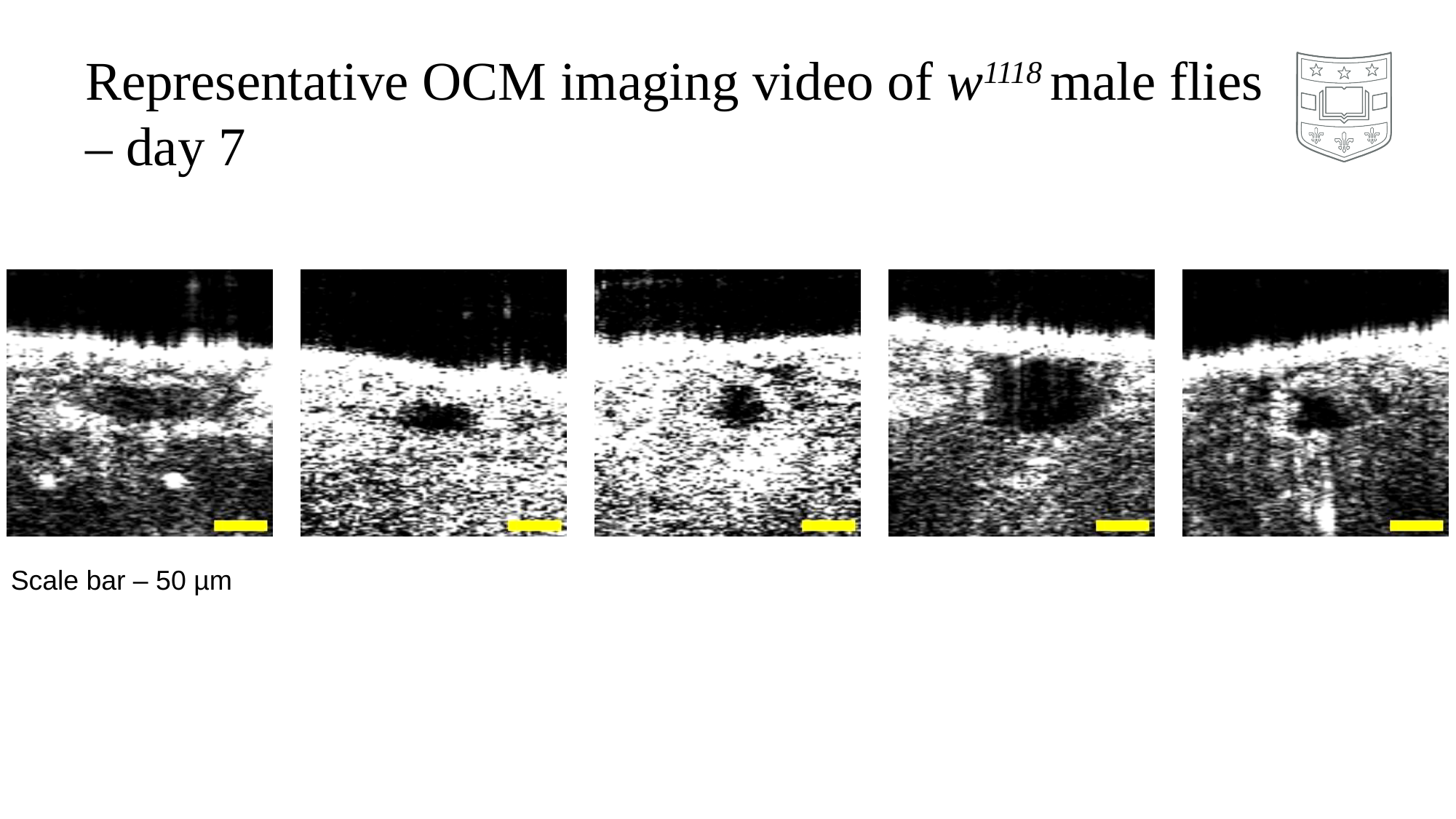

#### Representative OCM imaging video of w^1118^ male flies – day 7
Scale bar – 50 µm

### Slide 2
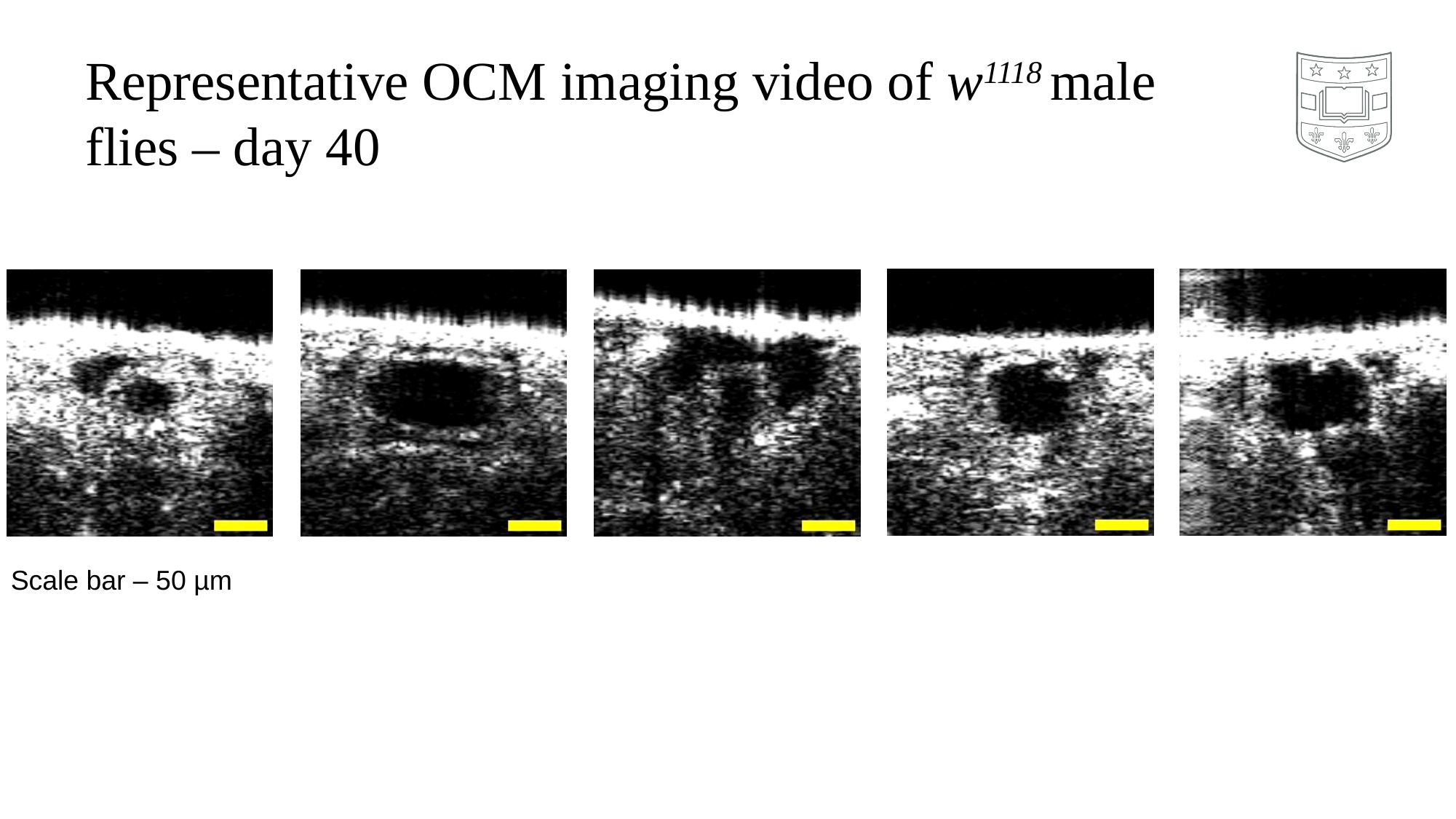

#### Representative OCM imaging video of w^1118^ male flies – day 40
Scale bar – 50 µm

### Slide 3
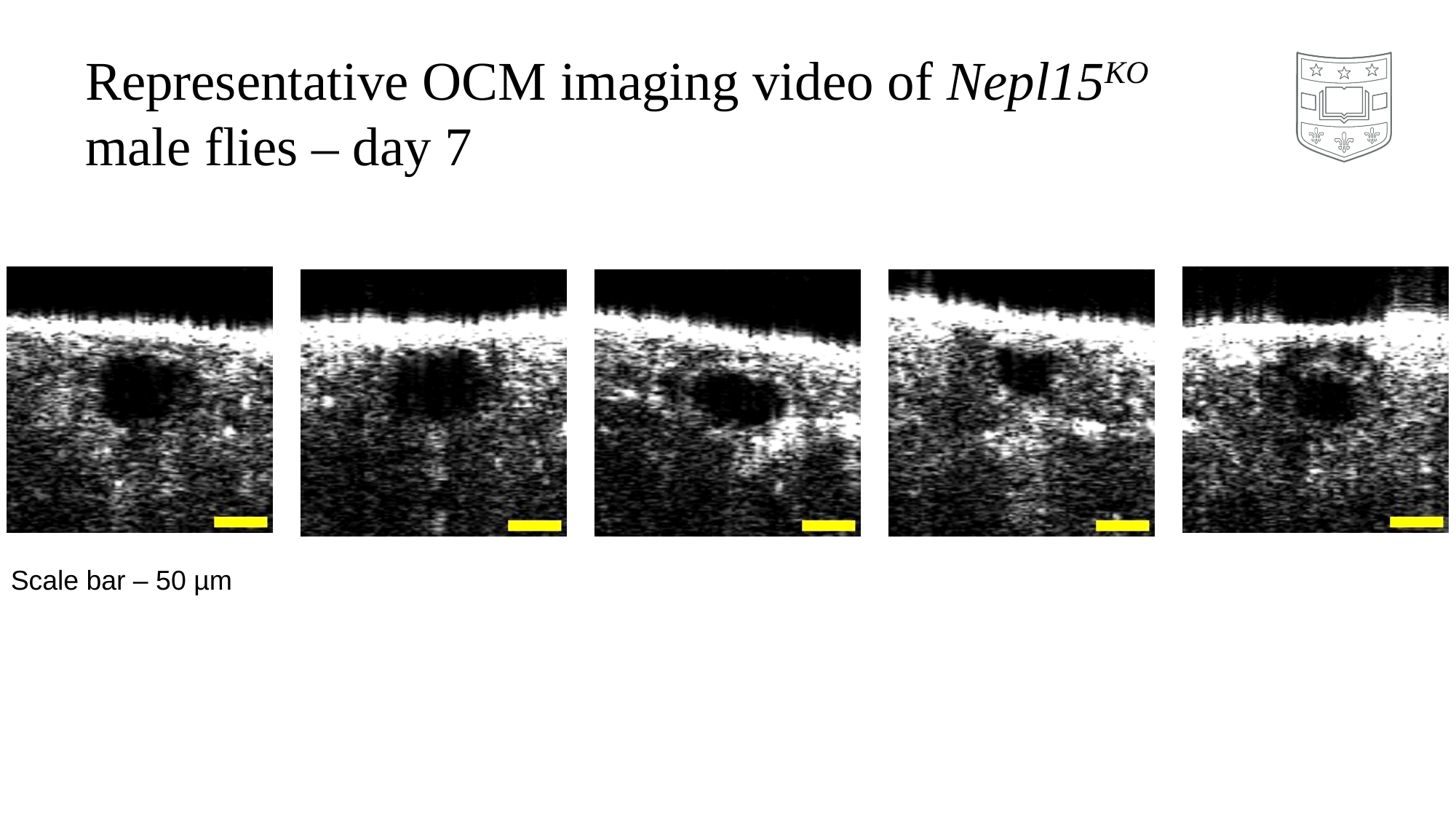

#### Representative OCM imaging video of Nepl15^KO^ male flies – day 7
Scale bar – 50 µm

### Slide 4
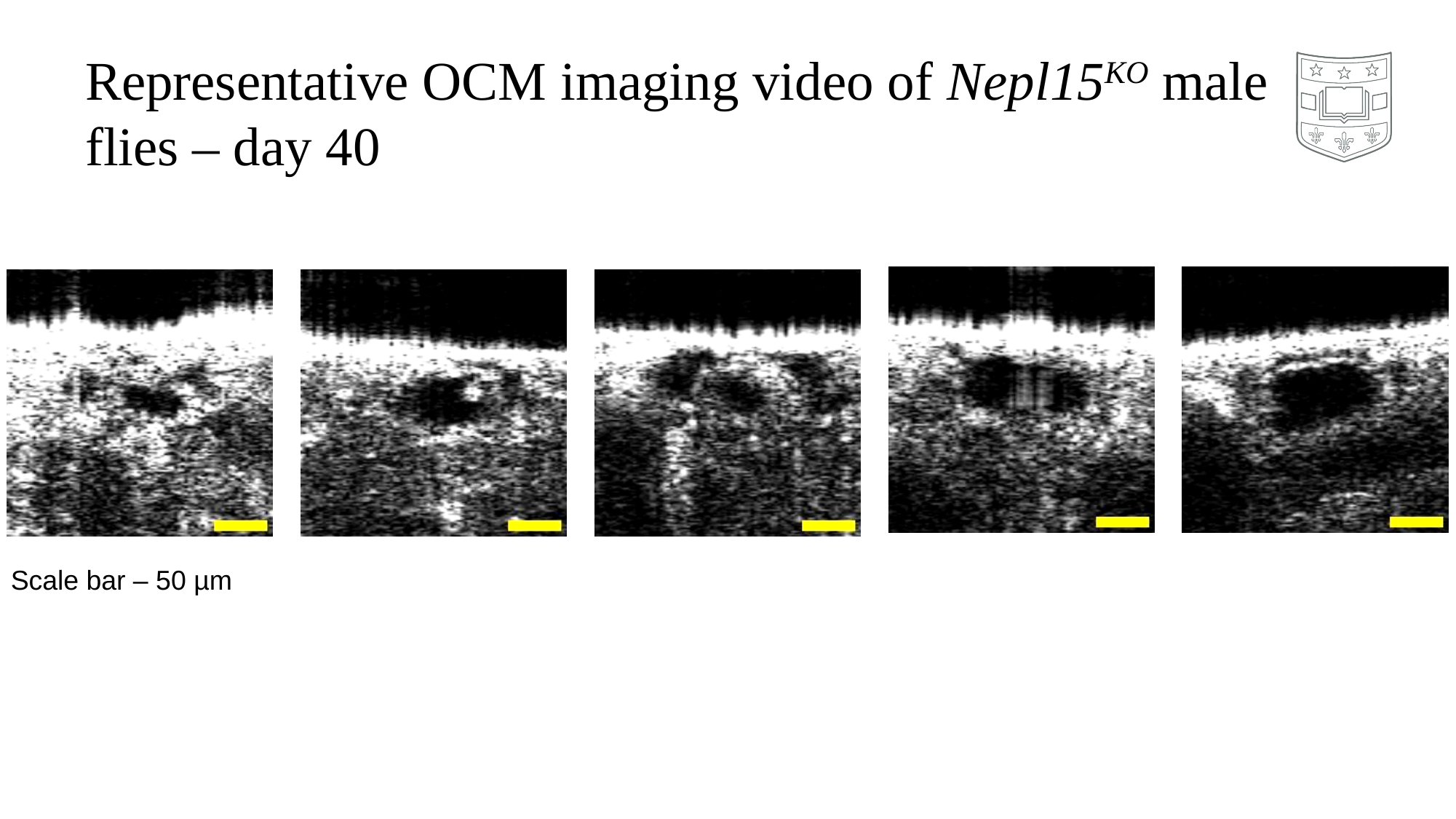

#### Representative OCM imaging video of Nepl15^KO^ male flies – day 40
Scale bar – 50 µm

### Slide 5
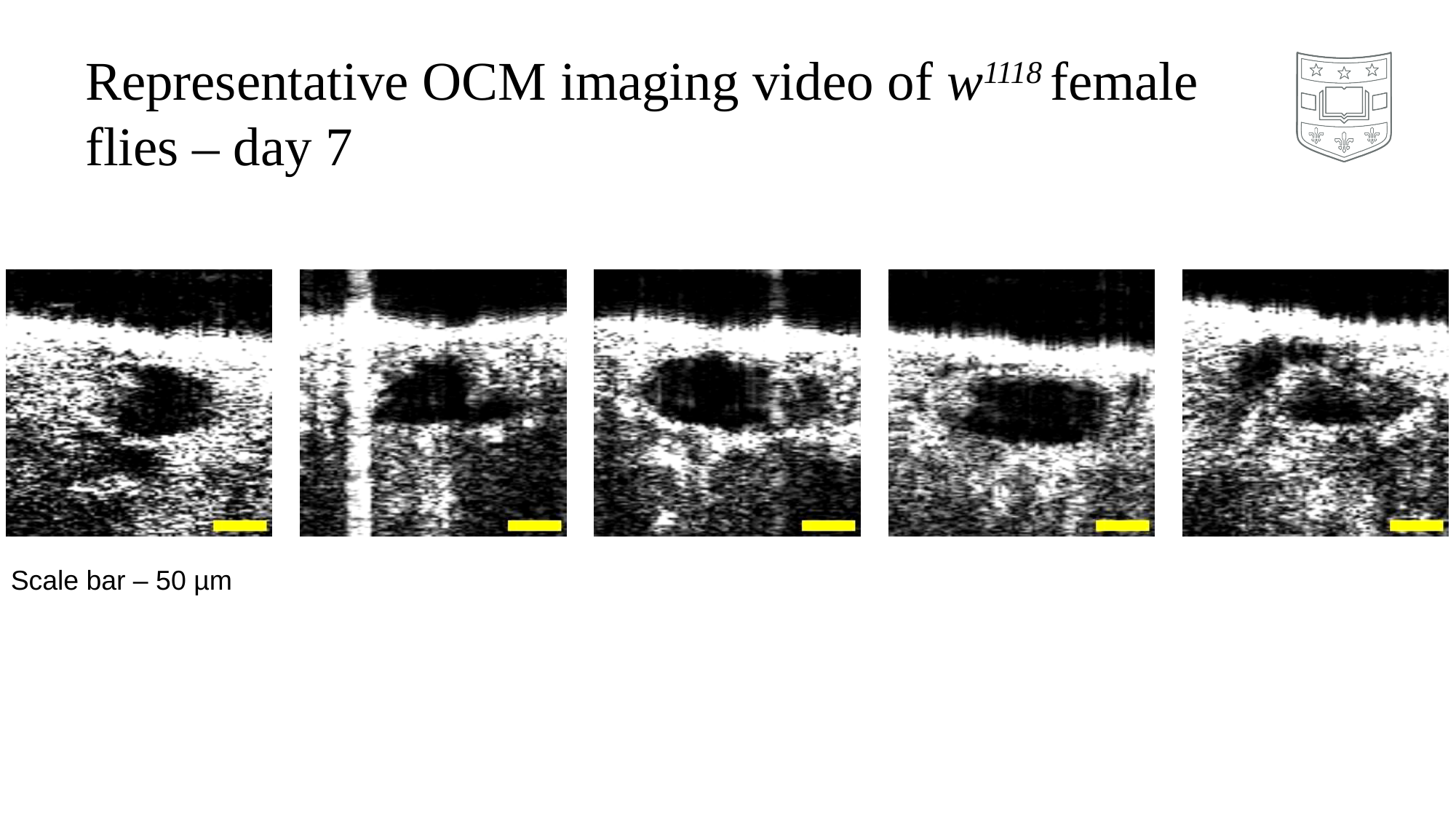

#### Representative OCM imaging video of w^1118^ female flies – day 7
Scale bar – 50 µm

### Slide 6
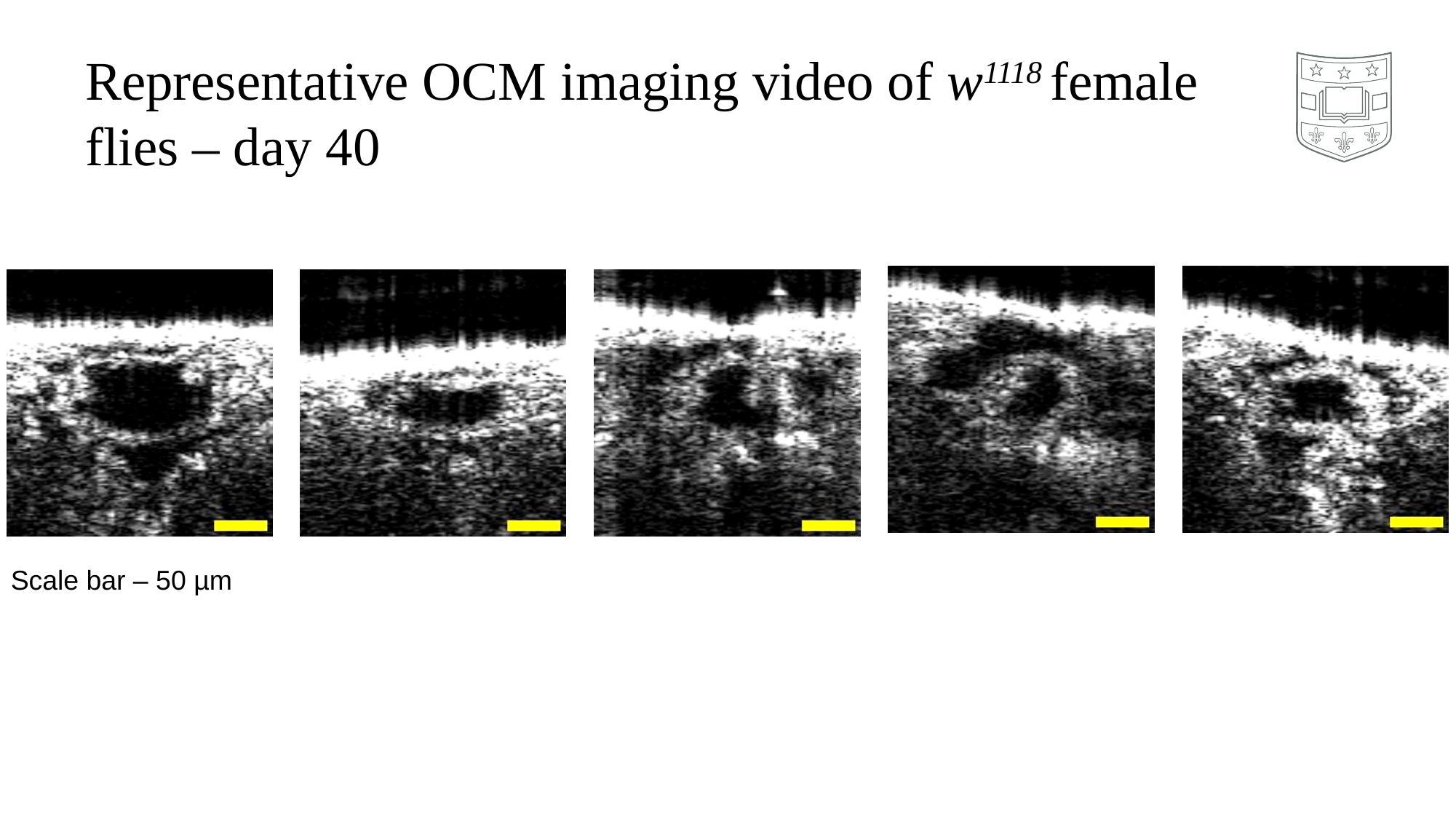

#### Representative OCM imaging video of w^1118^ female flies – day 40
Scale bar – 50 µm

### Slide 7
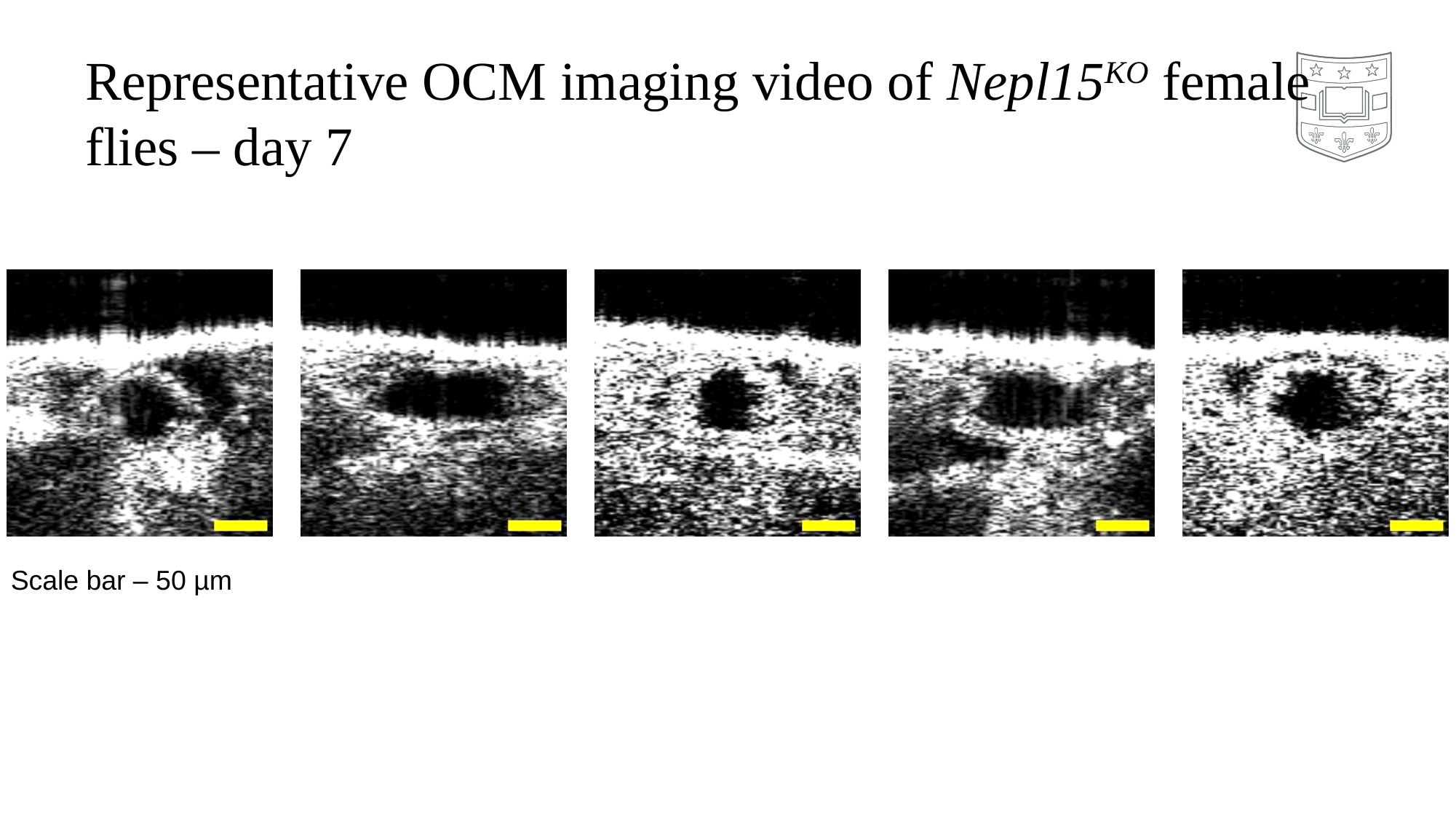

#### Representative OCM imaging video of Nepl15^KO^ female flies – day 7
Scale bar – 50 µm

### Slide 8
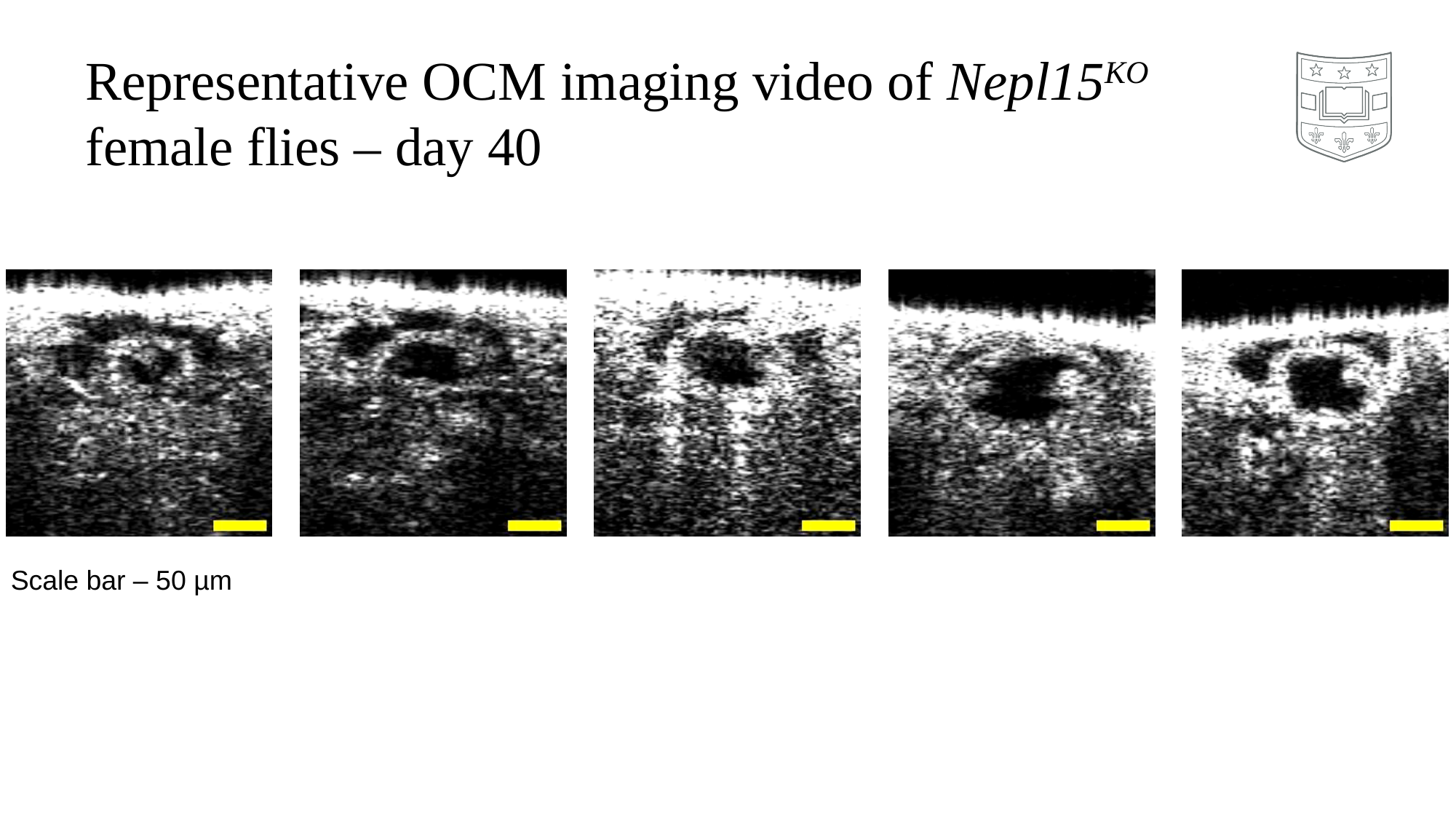

#### Representative OCM imaging video of Nepl15^KO^ female flies – day 40
Scale bar – 50 µm
